# Development of the axonal βII-spectrin periodic skeleton requires active cytoskeletal remodelling

**DOI:** 10.1101/2025.02.19.639207

**Authors:** Shivani Bodas, Ashish Mishra, Pramod Pullarkat, Aurnab Ghose

## Abstract

The axonal membrane-associated periodic skeleton (MPS), consisting of F-actin rings crosslinked by spectrin heterotetramers, is ubiquitous and critical for neuronal function and homoeostasis. However, the initiation and early development of the axonal MPS are poorly understood. Using superresolution imaging, we show that βII-spectrin is recruited early to the axonal cortex, followed by progressive establishment of long-range periodic order. Microtubule dynamics are essential for MPS formation in the early stages, but transition to a passive stabilising role in mature axons. We show that the early subplasmalemmal recruitment of βII-spectrin is dependent on cortical actin but not on actomyosin contractility, and active nucleation of F-actin is required in early development but is dispensable for the mature MPS. Using a βII-spectrin knockout model, we demonstrate that the actin-binding and lipid-interacting domains of βII-spectrin are critical for its subplasmalemmal confinement and, subsequently, MPS maturation. These findings highlight stage-specific cytoskeletal remodelling underlying MPS development and advance our understanding of axonal subcellular architecture.

## INTRODUCTION

The axonal subplasmalemmal membrane periodic skeleton (MPS) is ubiquitous across phyla and forms a quasi-1d lattice along the entire neuronal axon (D’Este et al., 2016; He et al., 2016). A similar scaffold is found in dendrites and dendritic spine necks (Han et al., 2017). The MPS comprises of axially aligned tetramers of α and β spectrins that are attached to evenly distributed radial F-actin rings, which maintain a typical spacing of 180 – 190 nm (Xu et al., 2013). The lattice acts as a load-bearing scaffold, providing mechanical support to the plasma membrane (Hammarlund et al., 2007; Dubey et al., 2020). The MPS plays an essential role in the maintenance of axonal homoeostasis, regulating the ion channel distribution, maintaining the axonal diameter and facilitating axonal transport (Costa *et al*., 2020a; Wang *et al*., 2020a).

Furthermore, the MPS acts as a signalling scaffold (Zhou et al., 2019), limits diffusion at the axon initial segment (Albrecht et al., 2016), modulates endocytosis (Wernert et al., 2024) and is linked to axon degeneration (Unsain et al., 2018; Wang et al., 2019). Studies in *C. elegans*, mice and humans have revealed that impairment of the MPS leads to a variety of defects and lethality(Hammarlund et al., 2007; Huang et al., 2017a; b; Lorenzo et al., 2019; Cousin et al., 2021).

Despite the explosion of interest in the axonal MPS, its early development and maintenance are not well understood. Cytoskeleton dependence on MPS stability has been typically studied in late-stage neurons and does not address early functions in initiating MPS development. The depolymerisation of F-actin inhibits MPS development in early axons (Xu et al., 2013; Zhong et al., 2014) and destabilises mature MPS (Qu et al., 2017; Unsain et al., 2018). However, what functions are subserved by F-actin in MPS initiation are unclear. Similarly, while microtubule integrity appears to be necessary for MPS maintenance (Zhong et al., 2014; Qu et al., 2017), its function in MPS development is unknown.

We present a chronological analysis of the βII-spectrin MPS development in embryonic DRG neurons using STED nanoscopy. Our findings reveal that βII-spectrin is recruited early to the axonal cortex and progressively matures to attain long-range periodic order and uniform spacing. In the early stages, microtubule dynamics are essential for MPS formation, but microtubules have a passive function in stabilising the mature MPS. We show that βII-spectrin is recruited to the axonal cortex via interactions with subplasmalemmal F-actin, and the early MPS is dependent on actively nucleating actin networks. However, the mature, stable MPS is insensitive to the inhibition of F-actin nucleation. Using βII-spectrin knockout cells, we demonstrate that the actin-binding and PH-domains of βII-spectrin are necessary for its recruitment to the axonal cortex.

## MATERIALS AND METHODS

### Primary neuronal cultures

All experimental procedures were conducted in accordance with guidelines approved by the Institutional Animal Ethics Committee of the Indian Institute of Science Education and Research (IISER) Pune, India. Fertilised White Leghorn chicken eggs were obtained from Venkateshwara Hatchery Private Limited (Pune, India) and were incubated at 37 °C in a humidified incubator for 8-9 days.

Dorsal root ganglia (DRG) were isolated from embryos using sterile Leibovitz’s L-15 medium (Hi-Media, AL204A) for dissection. Dissections were performed in a laminar flow hood. Around 12-16 DRGs were isolated per embryo in the L-15 medium. The suspension was centrifuged at 3000 rpm for 3 min and the supernatant was removed. The DRGs in the pellet were resuspended in trypsin solution and incubated for 30 min at 37 °C. This was followed by a 3-minute spin at 3000 rpm to remove most of the trypsin solution and an L-15 wash to remove any remaining trypsin. The dissociated neurons were then plated onto glass coverslips or Poly-D-lysine-coated (0.1 mg/ml PDL; Merck, A-003-E) glass-bottom dishes. Neurons were cultured in serum-free medium composed of L-15 supplemented with 6 mg/ml glucose, 1× GlutaMAX (Gibco, 35050061), 1× B27 supplement (Invitrogen, A1895601), 50 ng/ml nerve growth factor (Gibco, 13290010), 0.006 g/ml methyl cellulose (34516, ColorconID), and 1× PenStrep (HiMedia, A001A). Cultures were maintained in a non-CO2 incubator at 37 °C.

### Drug treatment of neurons

The specific drugs utilised, along with their corresponding treatment conditions, are described in the table below. All the cultures were fixed and processed for immunofluorescence analysis at different time points. All drugs were prepared in dimethyl sulfoxide (DMSO), and DMSO-treated cultures served as vehicle controls for each drug treatment.

| Drug | Source | Type of treatment | Time of addition | Concentration | Duration |
| --- | --- | --- | --- | --- | --- |
| Taxol | Santa Cruz (sc-201439) | Acute | DIV 1,2 & 4 | 10 µM | 30 min |
| Nocodazole | Tocris Bioscience (2792) | Acute | DIV 1,2 & 4 | 20 µM | 30 min |
| Latrunculin-A | Sigma Aldrich (L5163) | Acute | DIV 1,2 & 4 | 5 µM | 15 min |
| Jasplakinolide | Tocris Bioscience (1228) | Acute | DIV 1,2 & 4 | 5 µM | 30 min |
| Taxol | Santa Cruz (sc-201439) | Chronic | 16 hrs post seeding | 1 nM | Till DIV 2 or 4 |
| Nocodazole | Tocris Bioscience (2792) | Chronic | 16 hrs post seeding | 10 nM | Till DIV 2 or 4 |
| Jasplakinolide | Tocris Bioscience (1228) | Chronic | 16 hrs post seeding | 10 nM | Till DIV 2 or 4 |
| Blebbistatin | Sigma Aldrich (B0560) | Chronic | 16 hrs post seeding | 10 µM | Till DIV 2 or 4 |
| CK666 | Merck (182515) | Acute | DIV 2,4 & 6 | 20 µM | 30 min |
| SMIFH2 | Merck<br>(344092) | Acute | DIV 2,4 & 6 | 20 μM | 30 min |

### Cell fixation and immunostaining for fluorescence imaging

Cultured neurons were fixed in 4% (w/v) paraformaldehyde (PFA) in phosphate-buffered saline (PBS) for 10 min at room temperature (RT). PFA was removed by washing with PBS, and the cells were then permeabilised with 0.1% (v/v) Triton X-100 in PBS for 5 min at RT, followed by blocking in 3% (w/v) bovine serum albumin (BSA) in PBS for 1 hr at RT. The cells were then incubated with the primary antibodies – (βII-spectrin 1:700 (BD Biosciences; 612563), Anti-GFP 1:500 (Invitrogen; A11122) and βIII-tubulin 1:1000 (Santa-Cruz; sc-80005)) - overnight at 4°C. The primary antibodies were removed the following day, followed by 3x PBS washes, and the cells were incubated with the secondary antibodies – Alexa 488 conjugated anti-mouse or anti-rabbit (Invitrogen; A-11001, A-11008, respectively) – at 1:1000 dilution in blocking buffer for 1 hr at RT. For super-resolution microscopy, neurons at different stages of *in vitro* culture (days in vitro or DIV) were mounted in a mixture of 10% Mowiol 4–88 in poly (vinyl alcohol) (Sigma-Aldrich, 81381) and 2.5% (w/v) DABCO (1,4-diazabicyclo [2.2.2] octane; Sigma-Aldrich, D27802). The mounted samples were stored overnight in the dark at 4 °C before imaging.

### Plasmid construction and transfection

Human βII-Spectrin (from Addgene plasmid 31070: βII-Spectrin-HA) was incorporated into a pCAG-GFP containing vector by PCR. For the actin-binding deletion mutant (pCAG-ΔABD-βII-Spectrin-GFP) initial 3-277 aa was deleted from the pCAG-βII-Spectrin-GFP construct by PCR. Plasma membrane binding mutant (pCAG-βII-Spectrin-K2207Q-GFP) and Ankyrin binding mutant (pCAG-βII-Spectrin-Y187A-GFP) were created by introducing point mutations in the pCAG-βII-Spectrin-GFP construct by PCR. Cells were suspended in Optimem® (Gibco) medium with 15-20 μg of plasmid. Subsequently, the cell suspension was electroporated using the Nepagene NEPA21 Type II machine. The following parameters were used in all experiments: Poring pulse: 2x of 5 msec 125 V pulses with pulse interval of 50 msec and decay rate of 10%; polarity +. Transfer pulse: 5x of 50 msec 20 V pulses with pulse interval of 50 msec and decay rate of 40%; polarity +/-. After electroporation, the cells were seeded in glass-bottom dishes and incubated at 37 °C for relevant periods.

### βII-Spectrin intensity analysis

Axons from DIV 1, 2, and 4 were stained for βII-Spectrin and imaged using confocal microscopy. The middle region of the axon was selected and processed with ImageJ software for Otsu-based thresholding. A particle analyser tool was used to estimate the integrated density and area of the axon, followed by CTCF calculation.

### Fluorescence recovery after photobleaching & analysis

Primary neurons and differentiated βII-Spectrin KO SH-SY5Y cells were transfected via electroporation and used in experiments at various time points, ranging from 24 to 144 hr post-transfection. The following plasmids were used for transfection: pCAG-βII-Spectrin-GFP, pCAG-βII-Spectrin-K2207Q-GFP, pCAG-βII-Spectrin-Y187A-GFP, and pCAG-ΔABD-βII-spectrin-GFP.

Fluorescence recovery after photobleaching (FRAP) experiments were conducted using a Zeiss LSM 710 confocal microscope equipped with a 488 nm Argon laser for bleaching and a 37 °C humidified stage. Time-lapse images of GFP fluorescence were acquired at 2-second intervals using a Zeiss Plan-Apochromatic 63x/1.4 Oil objective with 3× optical zoom. Regions of interest (ROI) measuring a maximum of ∼2 × 1 microns (height × width) along the middle of the axon were iteratively photo-bleached (50-100 iterations) by scanning with a 488 nm laser at 100% power and a scanning speed of 6 μs/pixel. Post-bleach images were acquired with 15–30% laser intensity for 150 frames (2 second interval). The resulting images were analysed using the ImageJ software.

For each region of interest (ROI), background fluorescence was subtracted at each time point. A reference region was used to correct for fluorescence loss due to bleaching during image acquisition by normalising it to the pre-bleaching intensity. The background-corrected and acquisition-loss-corrected ROI intensities were then further normalised by setting the pre-bleaching intensity to 100% and the immediate post-bleaching intensity to 0%. The recovery curve was fitted to a single-phase association function from data plotted with a 2s or 10s interval. The extra sum-of-squares F-test was used to compare the plateaus of the curves. The half-life (t_½_) was estimated from the fitted curve of individual axons, and the % mobile fraction was calculated by averaging the last 5 frames of the recovering curve.

### SH-SY5Y Neuroblastoma cell differentiation and culture

SH-SY5Y cells were grown and maintained in 60 mm Petri dishes in standard culture medium (DMEM/F12 supplemented with 10% fetal bovine serum [FBS] and 1% penicillin-streptomycin) at 37 °C in a 5% CO2 atmosphere. Cells were passaged upon reaching 80–90% confluence using trypsin-EDTA, and all experiments were conducted using cells with a passage number below 20. SH-SY5Y cells were differentiated into neuron-like cells over 12-14 days of Retinoic acid (RA) and Brain-derived neurotrophic factor (BDNF) treatment. Briefly, cells were seeded in a surface-coated 60 mm flask and treated with retinoic acid in DMEM/F12 supplemented with 1% FBS for 5 days, followed by 8 days of brain-derived neurotrophic factor (BDNF) treatment in serum-free medium.

For FRAP and STED microscopy, cells were trypsinised, transfected with the appropriate plasmids, and then reseeded on glass-bottomed, surface-treated dishes after RA treatment. They were maintained in serum-free BDNF-supplemented media for 6 days, and experiments were performed.

### CRISPR/CAS9-mediated gene editing

Using an online CRISPR design tool (Ran et al., 2013), two pairs of oligo DNAs encoding gRNAs were designed to target βII-Spectrin exon 16 sequences (gRNA 1: 5′-CCAGACAGCGATCGCCTCGGAGG-3′ and gRNA 2: 5′-GAACGAGATCGACAACTACGAGG-3**′.** These oligos were synthesised, annealed, and inserted into the BbsI site of the pSpCas9(BB)-2A-GFP (PX458) plasmid (Addgene plasmid no.48138). The resulting plasmids were amplified and introduced into SH-SY5Y cells via electroporation.

GFP-positive cells were isolated through cell sorting. Western blot analysis was used to initially identify stably transfected clonal lines exhibiting complete depletion of SH-SY5Y expression.

### Indel analysis of βII-Spectrin knockout cell lines

To examine indel mutations in SH-SY5Y βII-Spectrin knockout cells, genomic DNA was extracted from the knockout clonal lines. Subsequently, PCR amplification of exon 16 was performed using the extracted genomic DNA as a template. The following oligonucleotides were used as primers: forward 5′-CCCTGAAAAACCGAGAGGCC-3′ and reverse 5′-GCTCGTTCCATCCAGTGTCC-3′. The amplified DNA was then purified, and the resulting plasmids were sequenced to identify indel mutations.

### STED imaging & analysis

STED imaging was conducted using a Leica TCS SP8 STED 3x microscope (Leica Microsystems) on medial axonal segments of embryonic chick DRG neurons and differentiated SH-SY5Y neurons. Images were captured with an HC PL APO CS2 100x/1.4 STED WHITE oil immersion objective (Leica Microsystems) in both confocal and STED modes. Acquisition settings included 16x line averaging and detector gating on a Hybrid Detector (HyD, Leica Microsystems) with a range of 0.3 ns to 8 ns. The zoom factor was set to ∼6x, with a pixel size of up to 20 nm. A 592 depletion laser was used for all image acquisitions. All other settings were kept constant.

Images were deconvolved using Huygens Professional software prior to analysis. Medial axonal regions were segmented into ∼3 µm sections, and intensity versus distance profiles were generated for each segment. Autocorrelation functions using MATLAB (R2021a) were computed based on the intensity profiles of each segment. To obtain the average autocorrelation function, autocorrelation curves from all the selected axon segments were averaged for each experimental condition. The autocorrelation amplitude was defined as the difference between the first peak and the average of two adjacent valleys in the averaged autocorrelation curve. The average periodicity (or spacing) of the membrane periodic skeleton (MPS) structure for each ∼3 μm segment was determined by the position of the first peak in the autocorrelation function. The diameter of each segment was calculated by measuring the edge-to-edge distance between spectrin signals.

### Axonal ablation and analysis

DRG neurons were grown on a glass-bottom dish without any coating to obtain detached axons. Axons of DIV1, DIV2, and DIV4 neurons were subjected to laser ablation using a home-built laser ablation setup. This setup consists of a 355 nm, 25 mJ, 350 ps pulsed laser (PowerChip PNV-M02510-100; Teem Photonics, Meylan, France) coupled to the side port of a Leica TCS SP8 confocal microscope using a custom filter-cube and focussed on the sample using a 40x/0.75 dry Phase Contrast objective. A custom-made 90° rotated filter cube having a UV-reflecting dichroic filter (T387lp-UF3, Chroma) was used to reflect the laser light into the objective. During and after ablation, phase contrast images were recorded using a CCD camera (Phantom VRI-MICRO-C210-16GB-M, Vision Research) at 2000 fps for 0.8 seconds to capture the initial fast retraction process of the ablated axons. A custom MATLAB code was employed to quantify the evolving gap distance. The x- and y-coordinates obtained were converted to distance values and plotted using GraphPad software.

Estimation statistics (www.estimationstats.com), Cohen’s d and a two-sided permutation test between the final gap length post-ablation were determined. The F-test was used to compare the plateaus of the evolving curve after fitting the data to a single-phase association model.

### Statistical analysis

All statistical analyses and visual representations were conducted using GraphPad Prism 10 (10.4.0) except for Fig. S1C. In the figures, data are presented with a central line indicating the mean, along with error bars displaying the standard error of the mean (SEM). The number of data points analysed for each graph is indicated in the figure legends. Statistical tests, along with correction tests used for the analysis, are mentioned in the figure legend. Estimation statistics (www.estimationstats.com) were used to plot the data and to calculate Cohen’s d in Fig. S1C.

## RESULTS

### Axonal βII-spectrin progressively develops a highly periodic organisation

To investigate the developmental trajectory of the spectrin MPS in embryonic DRG neurons, dissociated DRG neuron primary cultures were fixed at various time points, immunolabelled with βII-spectrin antibodies, and imaged using STED nanoscopy.

We quantified the early development of the MPS by imaging the medial segments of axons cultured for 1, 2, 4, and 6 days *in vitro* (DIV), using autocorrelation analysis of the βII-spectrin signal (Fig. 1A,B). We quantified the degree of periodicity (autocorrelation amplitude at the first peak) and the periodic spacing of the βII-spectrin rings (periodicity) from the autocorrelation curves (Fig. 1C,D). The representative micrographs (Fig. 1A) depict the emergence and evolution of the periodic organisation of βII-spectrin in the axons across development. βII-spectrin was already localised to the axonal shaft by DIV1, but its organisation was weakly periodic (Fig. 1A). However, the average autocorrelation amplitudes increased progressively with DIV (Fig. 1B,C), indicating the consolidation of long-range periodic order. Concomitant with the increase in the extent of periodicity, the spacing (periodicity) of the βII-spectrin signal (Fig. 1D) reduced with the age of the cultured neurons from 214 ± 6.17 nm in DIV1 to 196.7 ± 2.52 nm in DIV2. It further decreased to 192.9 ± 8.9 nm and 185.8 ± 4.3 nm on DIV 4 and DIV6, respectively. The developmental maturation of the MPS was further revealed by the progressive reduction of the coefficient of variation of the βII-spectrin spacing (Fig. 1D), decreasing from 25.9 at DIV1 to 4.3 at DIV6. The high variability of the periodicities during the early stages was expectedly associated with lower autocorrelation amplitudes (Fig. 1E). The autocorrelation amplitude increased with development as the spacing converged to 185.8 ± 4.3 nm by DIV 6. Interestingly, in these axonal segments, the amount of βII-spectrin is unchanged across DIV1, 2 and 4 (Fig. S1A), though the distribution becomes highly periodic. This suggests MPS maturation is dominated by the reorganisation of spectrin, recruited to the cortex early in development, from a relatively disordered to an ordered periodic scaffold.

**Figure 1:**
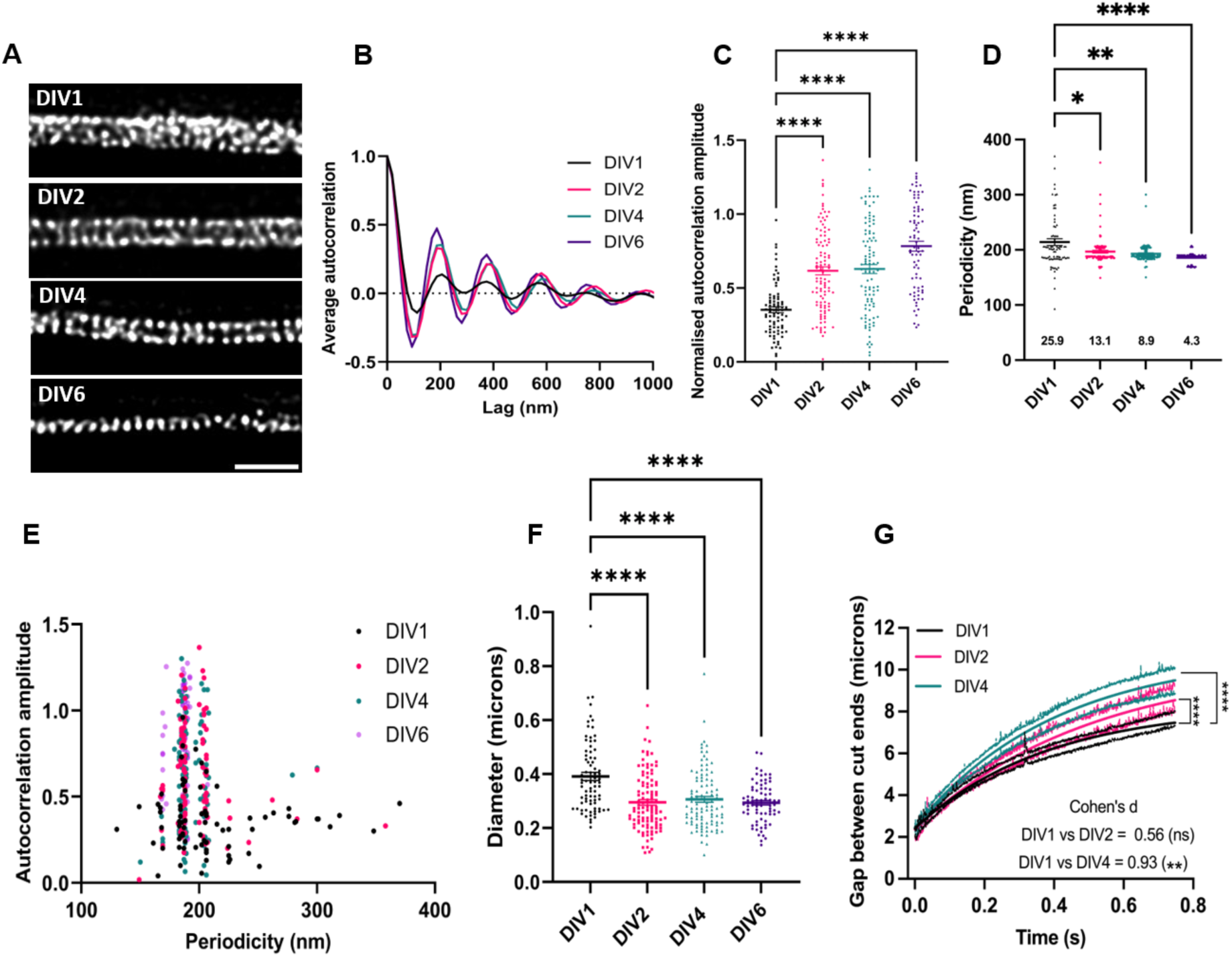
The βII-spectrin membrane periodic skeleton (MPS) assembles progressively. (A) Representative micrographs of single optical sections from STED nanoscopy of DIV1, DIV2, DIV4, and DIV6 DRG axons stained with βII-spectrin antibody. Scale bar: 1 µm. (B) Average autocorrelation curves across multiple axonal regions. (C) The normalised autocorrelation amplitudes of the βII-spectrin rings. Amplitudes were quantified as the difference between the first peak and the average of the first two valleys of the autocorrelation curve. The data were analysed using the Brown-Forsythe and Welch ANOVA test with multiple comparisons corrected by Games-Howell’s test. (D) Periodicity across DIVs demonstrates a reduction in the spacing between the βII-spectrin rings. The data were analysed using the Brown-Forsythe and Welch ANOVA test with multiple comparisons corrected by Games-Howell’s test. The numbers below each data set indicate the coefficient of variation of the distribution. (E) Plot of the normalised autocorrelation amplitudes against periodicity for all DIVs, indicating the degree of autocorrelation and the relative distribution of MPS periodicity. (F) βII-spectrin ring diameter across DIVs. The data were analysed using the Kruskal-Wallis test with multiple comparisons corrected by Dunn’s test. For A-F, 80-100 axonal segments from 3 biological replicates for each DIV were analysed. (G) The evolving gap length between cut ends following ablation is represented as a one-phase fit of the means (± SEM). Curve fits were compared using the extra sum-of-squares F-test. The effect sizes and significance, represented by Cohen’s d and two-sided t-test, obtained by comparing the gap length in the last 10 frames (till 0.74 sec) using Estimation statistics (Fig S1C) are indicated. The data are derived from n=19 (DIV1), n=15 (DIV2) and n=16 (DIV4) axons from at least three biological replicates. (ns, p > 0.05; *, p ≤ 0.05; **, p ≤ 0.01; ****, p ≤ 0.0001).

To evaluate whether MPS maturation across these early developmental stages influenced the axonal morphology, we measured the diameter of the βII-spectrin rings as a surrogate measure of axonal diameter. MPS maturation across DIVs correlated with a reduction in diameter (Fig.1F). This observation is consistent with the role of the MPS in regulating the axonal diameter (Costa et al., 2020; Wang et al., 2020) and suggests progressive constriction of the axonal shaft as the MPS matures. Interestingly, the decrease in diameter was already substantial by DIV2. This observation, together with a significant increase in autocorrelation amplitude (Fig. 1B) in DIV2 (compared to DIV1), suggests an early maturation of the βII-spectrin MPS in DRG neurons.

The axon is actively contractile (Mutalik et al., 2018), and the MPS has been associated with axonal tension and pre-stress (Krieg et al., 2014; Dubey et al., 2020). To assess if early MPS maturation contributes to axonal tension, we employed laser-induced axonal ablation of DIV1, 2 and 4 neurons (Fig. S1B) and followed the evolution of the gap distance between the two ends following ablation. The rate of change of the gap distance may depend on passive stresses, active contractility or progressive cytoskeletal remodelling. However, the early, exponential response is likely to be dominated by axonal pre-tension. F-test comparing the plateaus of the one-phase association fits revealed that DIV4 axons exhibit greater pre-stress than DIV1 and DIV2 axons (Fig. 1G), while the quantitative analysis of gap distance at 0.74 seconds revealed a substantial effect size (Cohen’s d=0.93; p=0.0078) compared to DIV1 (Figs. 1G, S1C). Maturation of the βII-spectrin MPS thus appears to correlate with the evolution of axonal tension. However, the change observed is modest, and there is substantial pre-tension at DIV1 when the βII-spectrin MPS is only weakly periodic.

Collectively, these data reveal the maturation kinetics of the βII-spectrin MPS in DRG axons in early development. βII-spectrin is recruited early to the axonal cortex, followed by progressive establishment of periodic order, which is concomitant with a decrease in axonal diameter and an increase in axonal pre-stress.

### F-actin and microtubules are required for βII-spectrin MPS development

Several studies have implicated both the actin and microtubule cytoskeletal systems in the maintenance of the F-actin/βII-spectrin MPS, with most focussing on late-stage neurons. We evaluated the role of axonal F-actin and microtubules in the early development of the βII-spectrin MPS. DRG neurons at DIV1, 2, and 4 were acutely exposed to drugs that destabilised or stabilised the two cytoskeletal systems before fixation and evaluation of the βII-spectrin MPS via STED nanoscopy.

Suppression of F-actin polymerisation by latrunculin-A (Lat-A; 5 μM, 15 min) or microtubule polymerisation by nocodazole (Noco; 20 μM, 30 min) disrupted the long-range order of the periodic βII-spectrin and reduced the autocorrelation amplitudes across all DIVs (Fig. 2A-B,D-E,G-H). Furthermore, periodicity analysis revealed an increased coefficient of variation of the MPS spacing upon depolymerisation of the actin or microtubule cytoskeletons (Fig. 2C, F,I). In line with the involvement of the MPS in maintaining axon diameter, depolymerisation of F-actin or microtubules led to an increase in the diameter of the βII-spectrin rings at all DIVs (Fig. S2A, B, C).

**Figure 2:**
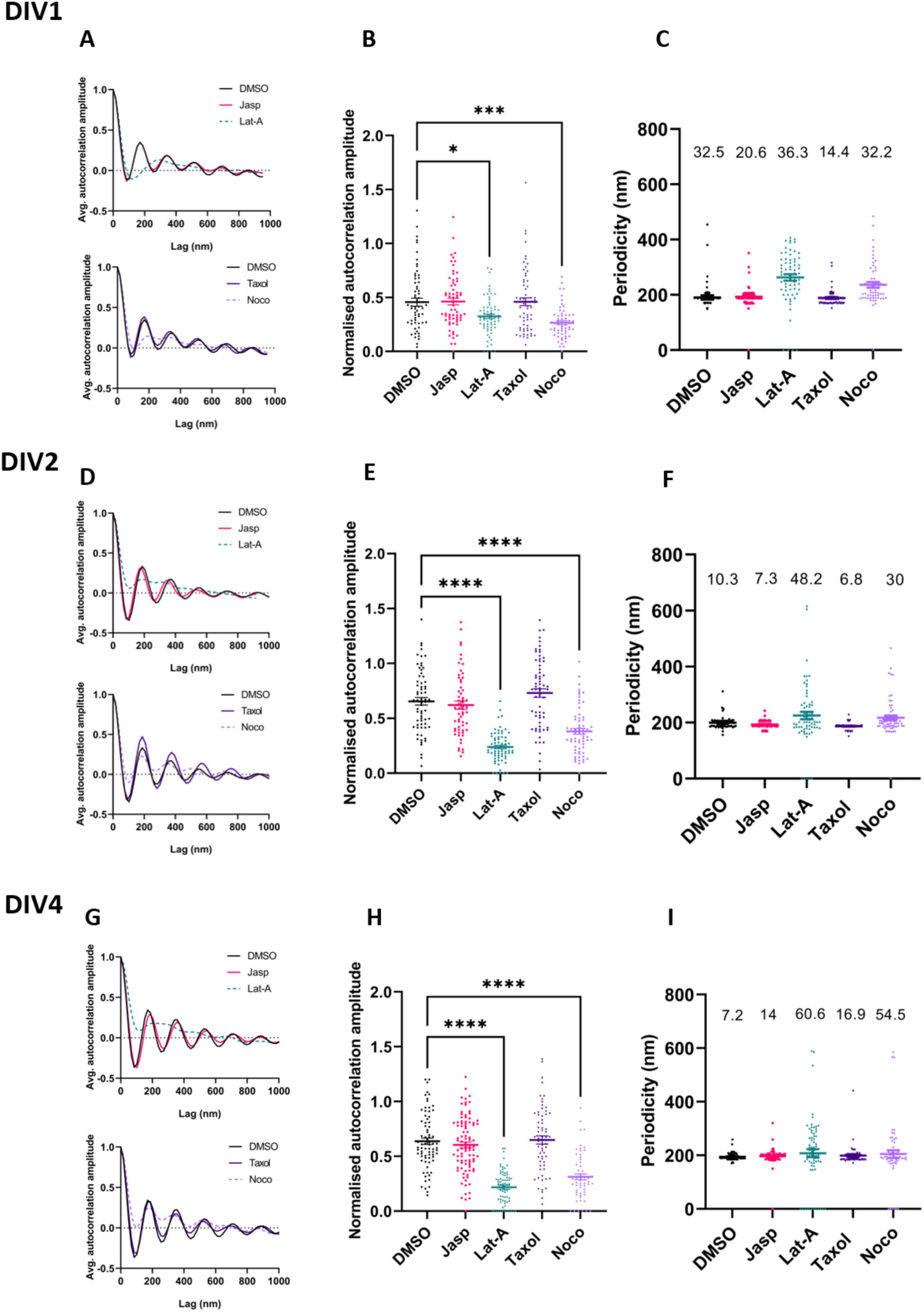
Destabilisation of F-actin and microtubules affects MPS development and stability. (A,D,G) Average autocorrelation curves of axonal segments across DIVs treated with DMSO (control), Jasplakinolide (Jasp), Latrunculin-A (Lat-A), Paclitaxel (Taxol), and Nocodazole (Noco). (B,E,H) The normalised autocorrelation amplitudes of the βII-spectrin rings across treatments and DIVs. The data were analysed using the Brown-Forsythe and Welch ANOVA test with multiple comparisons corrected by Games-Howell’s test (ns, p > 0.05; *, p ≤ 0.05; ***, p ≤ 0.001; ****, p ≤ 0.0001). (C, F, I) Periodicity (spacing) across DIVs treated with indicated drugs. The numbers below each data set indicate the coefficient of variation of the distribution. The numbers above each data set indicate the coefficient of variation of the distribution. For A-I, 60-95 axonal segments from 3 biological replicates for each DIV were analysed.

Acute treatment with the F-actin stabilising jasplakinolide (Jasp; 5 μM, 30 min) or microtubule stabilising paclitaxel (Taxol; 10 μM, 30 min), however, did not affect the autocorrelation or average autocorrelation amplitudes at any of the DIVs (Fig. 2A-B,D-E,G-H). While the periodicity analysis revealed no difference in the mean βII-spectrin spacing, there was a notable decrease in the coefficient of variation of the spacing upon either actin or microtubule stabilisation at DIV1 but it was not obvious at DIV2 and DIV4 when the MPS is already well organised (Fig. 2C,F,I). The latter result may suggest that pharmacological stabilisation of labile F-actin and microtubules in early developmental stages facilitates the maturation of the βII-spectrin MPS. The lability of F-actin at early stages of MPS development has been previously suggested as periodic F-actin is not detectable in young hippocampal axons, though the spectrin MPS is easily discernible (Zhong et al., 2014). Stabilisation of F-actin or microtubules did not change the βII-spectrin diameter at any DIV (Fig. S2A, B, C).

These experiments demonstrate the requirement of both F-actin and microtubules in the development of the βII-spectrin MPS.

### Microtubule dynamics are necessary to initiate MPS development

The involvement of microtubules in destabilising the early βII-spectrin MPS (Fig. 2) led us to investigate whether microtubule dynamics were required for the initiation and consolidation of MPS. We developed an early, chronic treatment protocol where a low dose of Noco (10 nM) or Taxol (1 nM) was added 16 h post-seeding (to allow for neuritogenesis) and the cultures fixed at DIV2 and DIV4 (Fig. 3A). Chronic disruption of microtubule polymerisation by Noco disrupted MPS formation at both DIV 2 and 4, as shown by the averaged autocorrelation curves and reduced autocorrelation amplitudes (Fig. 3B-D). Interestingly, and in contrast with acute exposure to Taxol (Fig. 2), chronic stabilisation of microtubules by Taxol compromised MPS formation (Fig. 3B-D) at both DIV2 and 4. Both microtubule depolymerisation and stabilisation increased the coefficient of variation of the periodicity (Fig. 3E). The compromised MPS formation was also reflected in diameter analysis, where both Nocodazole and Taxol treatments resulted in increased axonal diameter at both DIV2 and DIV4 (Fig. S3A). Notably, despite the compromised MPS formation, the amount of βII-spectrin in these medial axonal segments remained unchanged across DIV2 and DIV4 following chronic treatment with microtubule depolymerising or stabilising agents (Fig. S3B). This observation suggests that microtubules play a critical role in facilitating the organisational plasticity of βII-spectrin, rather than the recruitment of βII-spectrin to the axons.

**Figure 3:**
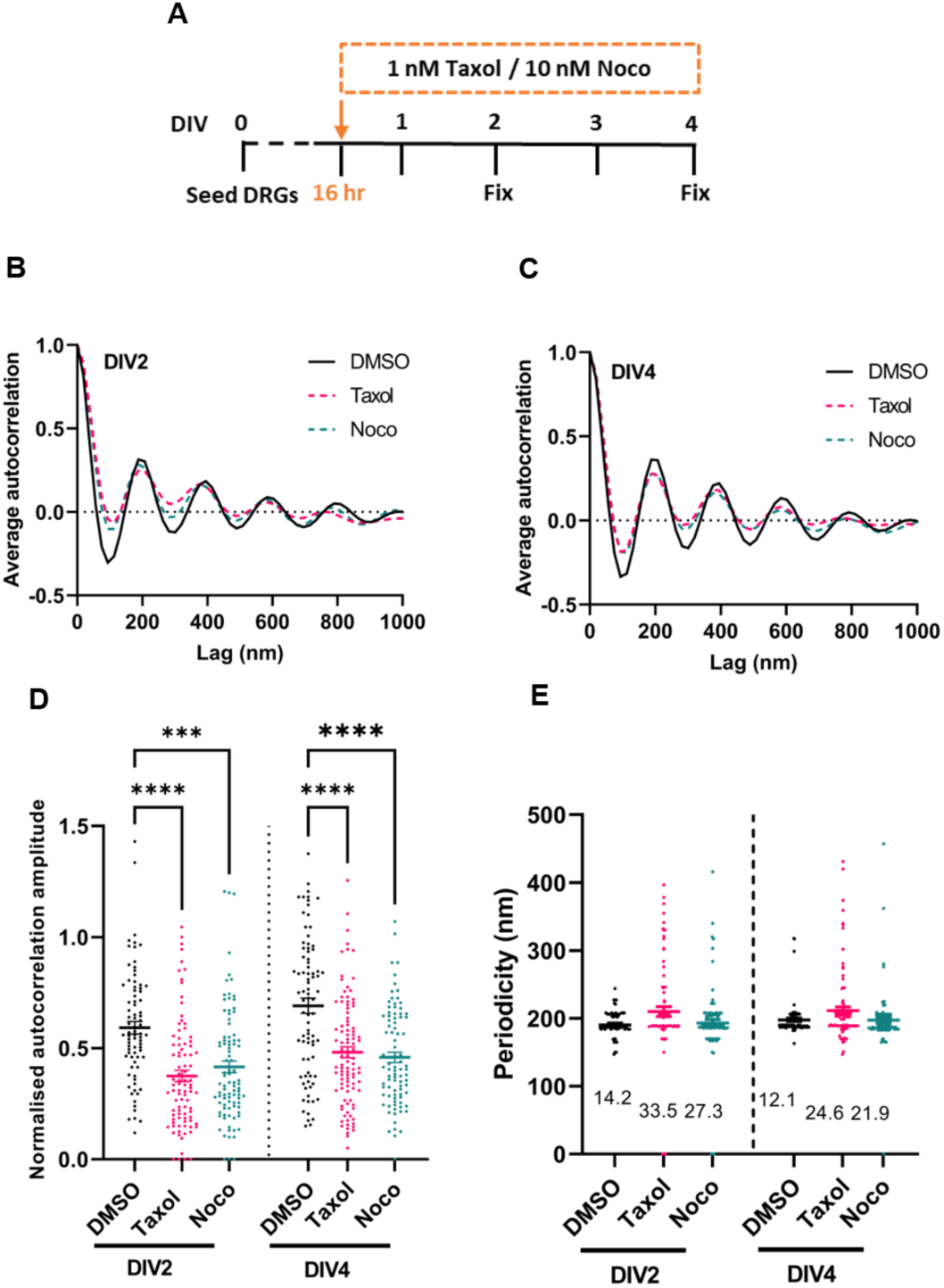
Microtubule dynamics are crucial for MPS formation. (A) Timeline of the experiment. 1 nM Taxol or 10 nM Nocodazole was added 16 hr after seeding the neurons and fixed either after another 32 hrs (DIV2) or after 80 hrs (DIV4). (B,C) Average autocorrelation curves drop for βII-spectrin stained axons from cultures treated with 10 nM Nocodazole (green dotted curve) and 1 nM Taxol (pink dotted curve) compared to DMSO (black curve). (D)The normalised autocorrelation amplitudes of the βII-spectrin rings. The data were analysed using the Brown-Forsythe and Welch ANOVA test with multiple comparisons corrected by Games-Howell’s test (***, p ≤ 0.001; ****, p ≤ 0.0001). (E) Periodicity across DIVs demonstrates an increase in variability between the βII-spectrin rings. The numbers below each data set indicate the coefficient of variation of the distribution. For B-E, 82-100 axonal segments from 3 biological replicates for each DIV were analysed.

Chronic treatment of hippocampal neurons with paclitaxel has previously been shown to induce multiple axon-like processes with MPS (Zhong et al., 2014). However, this study was performed at later stages, and the MPS of these axon-like processes were not quantitatively evaluated. Our data suggest that a balance of stable and dynamic microtubules is required to initiate MPS formation (Fig. 3), however, the maintenance of the relatively more established MPS is only sensitive to microtubule depolymerisation (Fig.2) but not stabilisation. At later stages, microtubules appear to have a more structural function in supporting the mature MPS, but microtubule dynamics in early stages seem to be involved in the generation of periodically organised βII-spectrin.

### F-actin stabilisation or actomyosin activity does affect axonal MPS development

A recent study in fibroblasts has suggested that acto-myosin contractility may drive spectrin clustering and periodic distribution of βII-spectrin (Ghisleni et al., 2024). Suppression of actin dynamics (treatment with jasplakinolide) increased periodic βII-spectrin clusters via myosin activity, while the inhibition of non-muscle myosin II (by blebbistatin) dissipated the βII-spectrin clusters (Ghisleni et al., 2024).

In axons, however, an acute reduction of F-actin turnover by jasplakinolide (Jasp) did not affect the MPS (Fig. 2). We tested if a chronic, low-dose stabilisation of F-actin would affect MPS development. 10 nM Jasp was added to DRG cultures 16 h post-seeding, and neurons were fixed at DIV2 and DIV4. As reported earlier, Jasp treatment resulted in shorter axons (Gallo et al., 2002), but the development of the βII-spectrin MPS was unaffected at both DIV2 and 4 (Fig. 4) compared to vehicle treatment. Curiously, the axons chronically exposed to Jasp were substantially thicker compared to the controls, and this was reflected in the increased βII-spectrin ring diameter (Fig. S3C). The diameter of the βII-spectrin rings did not change in response to acute treatment with Jasp as described earlier (Fig. S2), and it appears to be specific to long-term exposure and not dependent on the absence of the βII-spectrin MPS. In addition to stabilising F-actin, Jasp can also nucleate F-actin under certain conditions (Bubb et al., 2000). Chronic exposure to Jasp may lead to excessive, hyperstabilised axonal F-actin that results in short and thick axons. The stronger spectrin signal in Jasp-treated axon compared to controls observed may reflect the latter speculation (Fig. S3D).

**Figure 4:**
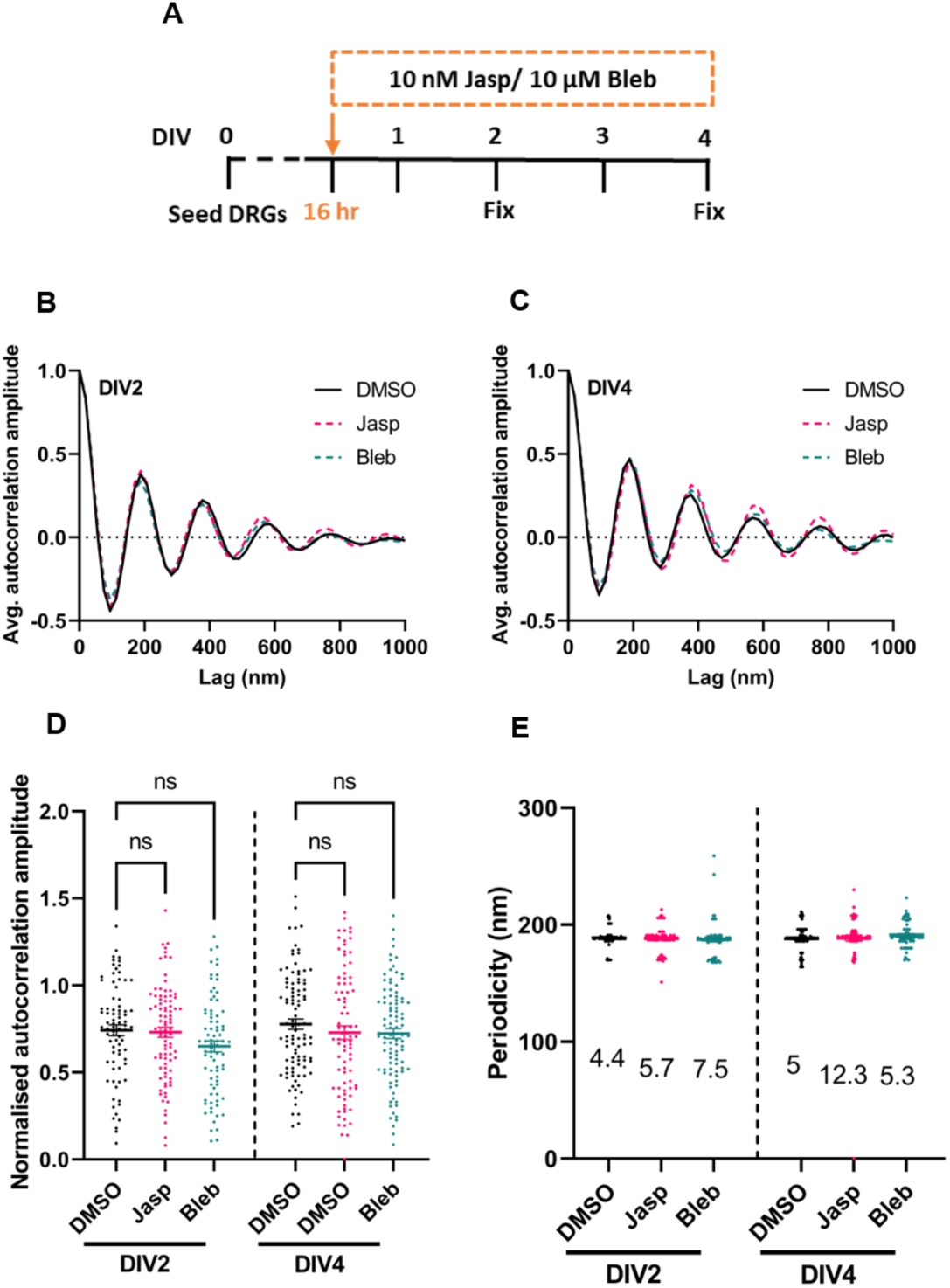
Actin stabilisation or myosin contractility does not affect MPS formation. (A) Timeline of the experiment. 10 nM Jasplakinolide or 10 μM Blebbistatin was added 16 hr after seeding the neurons and fixed either after another 32 hrs (DIV2) or after 80 hrs (DIV4). (B, C) Average autocorrelation curves show no difference for βII-spectrin stained axons from cultures treated with 10 μM Blebbistatin (green dotted curve) and 10 nM Jasplakinolide (pink dotted curve) compared to DMSO (black curve). (D)The normalised autocorrelation amplitudes of the βII-spectrin rings. The data were analysed using the Brown-Forsythe and Welch ANOVA test with multiple comparisons corrected by Games-Howell’s test (ns, p > 0.05). (E) Periodicity across DIVs demonstrates variability between the βII-spectrin rings. The numbers below each data set indicate the coefficient of variation of the distribution. For B-E, 79-98 axonal segments from 3 biological replicates for each DIV were analysed.

Acute inhibition of myosin activity has been previously shown to increase axonal and βII-spectrin ring diameter without affecting the periodic organisation of the MPS (Costa et al., 2020; Wang et al., 2020; Zhou et al., 2022). We again developed a low-dose, chronic exposure protocol with 10 μM blebbistatin (Bleb) added to DRG cultures 16 h post-seeding and the neurons fixed for analysis at DIV2 and DIV4. Low-dose, prolonged inhibition of non-muscle myosin II did affect the development of the βII-spectrin MPS (Fig. 4). However, consistent with the involvement of myosin contractility in maintaining axonal diameter (Costa et al., 2020; Wang et al., 2020), treatment with a low concentration of Bleb increased the average diameter of βII-spectrin rings compared to controls (Fig. S3C).

Our data suggest that, unlike in fibroblasts, actomyosin activity does not drive the assembly of periodic βII-spectrin clusters in axons.

### Actin nucleators Arp2/3 and Formins regulate MPS development in the early but not late stages

F-actin structures in cells are initiated by the activity of actin nucleators, which also modulate the morphology and dynamics of these structures. Previous reports indicate that MPS formation is impaired in neurons derived from *Drosophila* carrying loss-of-function mutations in the actin nucleators Arp2/3 and the formin DAAM (Qu et al., 2017). Given the dependence of the βII-spectrin MPS on F-actin (Fig. 2), we investigated the contribution of Arp2/3 and formins in MPS organisation at different developmental time points.

We exposed DRG neurons at different DIVs to the Arp2/3 complex inhibitor, CK666 (20 µM for 30 mins) or the formin inhibitor, SMIFH2 (20 µM for 30 mins). Inhibition of Arp2/3 or the formins in early-stage neurons (DIV2 and DIV4) led to the disorganisation of the βII-spectrin MPS with decreased autocorrelation and average autocorrelation amplitudes (Fig. 5A-C). These were accompanied by an increase in the variability of the MPS spacing, though the mean values were not statistically different. However, at DIV 6, inhibition of neither the Arp2/3 complex nor formin activity disrupted the abundance or periodicity of the MPS. These results suggest that MPS development requires both the linear (formins) and branched nucleators (Arp2/3) at the early developmental stages; however, actin nucleation is dispensable for the maintenance of stable MPS.

**Figure 5:**
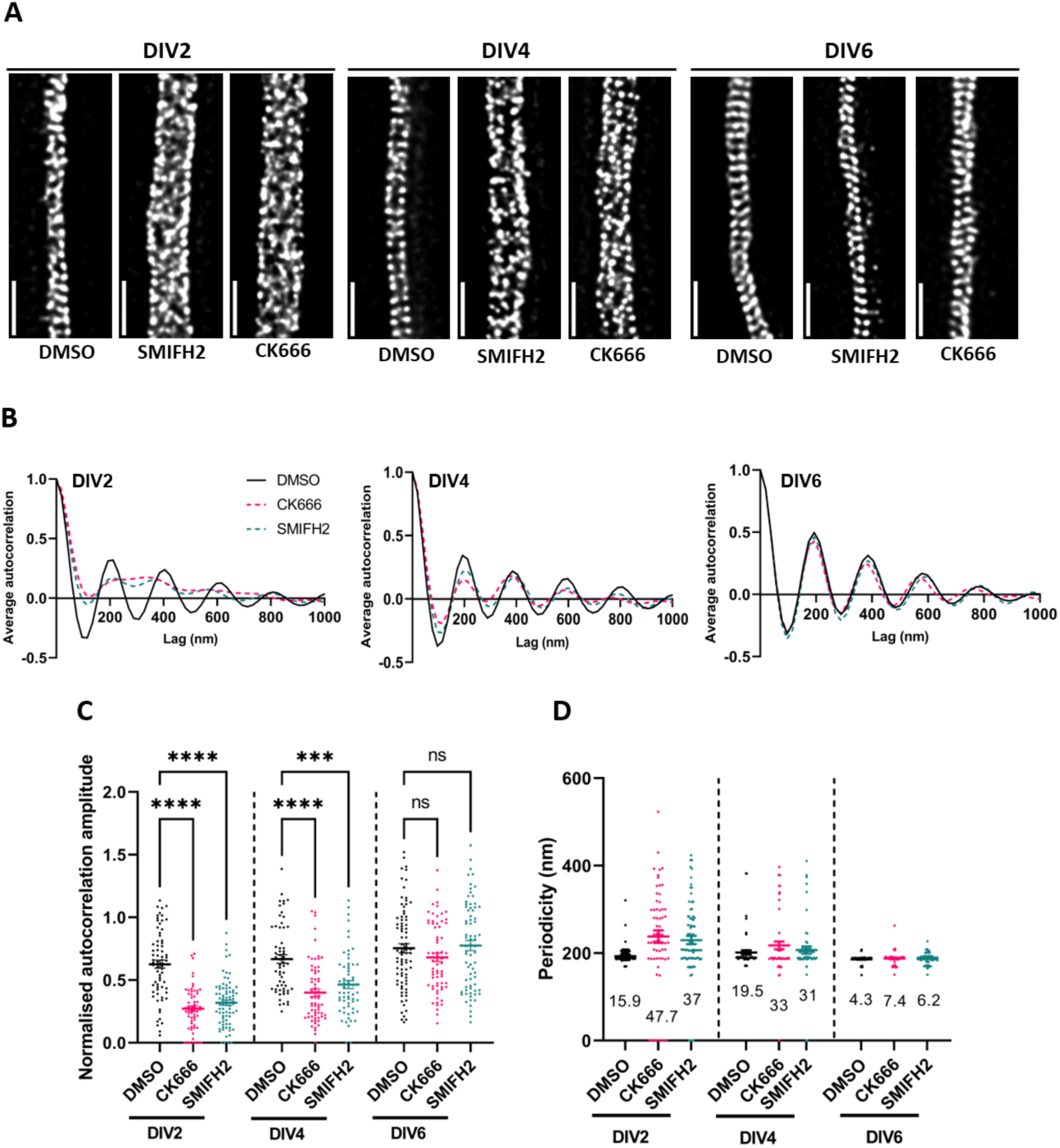
Formins and Arp2/3 affect MPS formation and maintenance. (A) Representative micrographs of single optical sections from STED nanoscopy of DIV2, DIV4, and DIV6 DRG axons stained with βII-spectrin post-treatment with DMSO control, Formin inhibitor (SMIFH2; 20 µM, 30 min) or Arp2/3 inhibitor (CK666; 20 µM, 30 min). Scale bar: 1 µm (B) Average autocorrelation curves across multiple axonal regions for all treatments. (C) The normalised autocorrelation amplitudes of the βII-spectrin rings. The data were analysed using the Brown-Forsythe and Welch ANOVA test with multiple comparisons corrected by Games -Howell’s test (ns, p > 0.05; ***, p ≤ 0.001, ****, p ≤ 0.0001) (D) Periodicity across DIVs demonstrates an increase in variability between the βII-spectrin rings at DIV2 and 4. The numbers below each data set indicate the coefficient of variation of the distribution. For B-D, 60-70 axonal segments from 3 biological replicates for each DIV were analysed.

The formin inhibitor, SMIFH2, also inhibits myosin II at high doses (Nishimura et al., 2021). As inhibition of myosin II increases the axonal diameter (Costa et al., 2020; Wang et al., 2020), we tested if treatment with SMIFH2 altered axonal diameter at DIV6 (no effect on MPS organisation at this stage). No change in the diameter of the βII-spectrin ring was observed following SMIFH2 treatment of DIV6 neurons (Fig S5), suggesting that the dose used in our experiments does not inhibit myosin II significantly. However, the compromised periodic organisation of βII-spectrin observed in early DIV2 and 4 upon treatment with either SMIFH2 or CK666 resulted in increased βII-spectrin ring diameter only at DIV2 and DIV4 but not at DIV6 (Fig. S5).

F-actin is necessary to crosslink multiple spectrin heterotetramers to form the mature axially concatenated arrangement of alternating F-actin rings and spectrins. In addition, the recruitment and retention of spectrin heterotetramers to the axonal cortex, prior to the development of a strongly periodic scaffold, could depend on subplasmalemmal F-actin. Our data indicating the sensitivity of the earlier stages to the inhibition of F-actin nucleation suggest labile F-actin structures with faster F-actin turnover at these stages. However, at later stages (DIV6), F-actin structures are more stable with reduced dependency on fresh nucleation and/or formin-dependent filament elongation.

### βII-spectrin progressively associates with the axonal cortex via interactions with F-actin

As the axon develops, βII-spectrin is recruited to the cellular cortex and progressively evolves into a highly periodic organisation (Fig. 1). We find early subplasmalemmal recruitment of βII-spectrin by DIV1 (Fig. 1A-D) and a dependence on F-actin (Figs. 2 and 5) in initiating MPS formation. To evaluate the dynamics of subplasmalemmal βII-spectrin, we employed fluorescence recovery after photobleaching (FRAP) analysis of DRG neurons transfected with GFP-tagged βII spectrin (βII-spectrin-GFP) at different developmental stages (Fig. 6A). To exclude cytoplasmic spectrin resulting from overexpression, only axons with low expression of βII spectrin-GFP were selected for the analysis.

**Figure 6:**
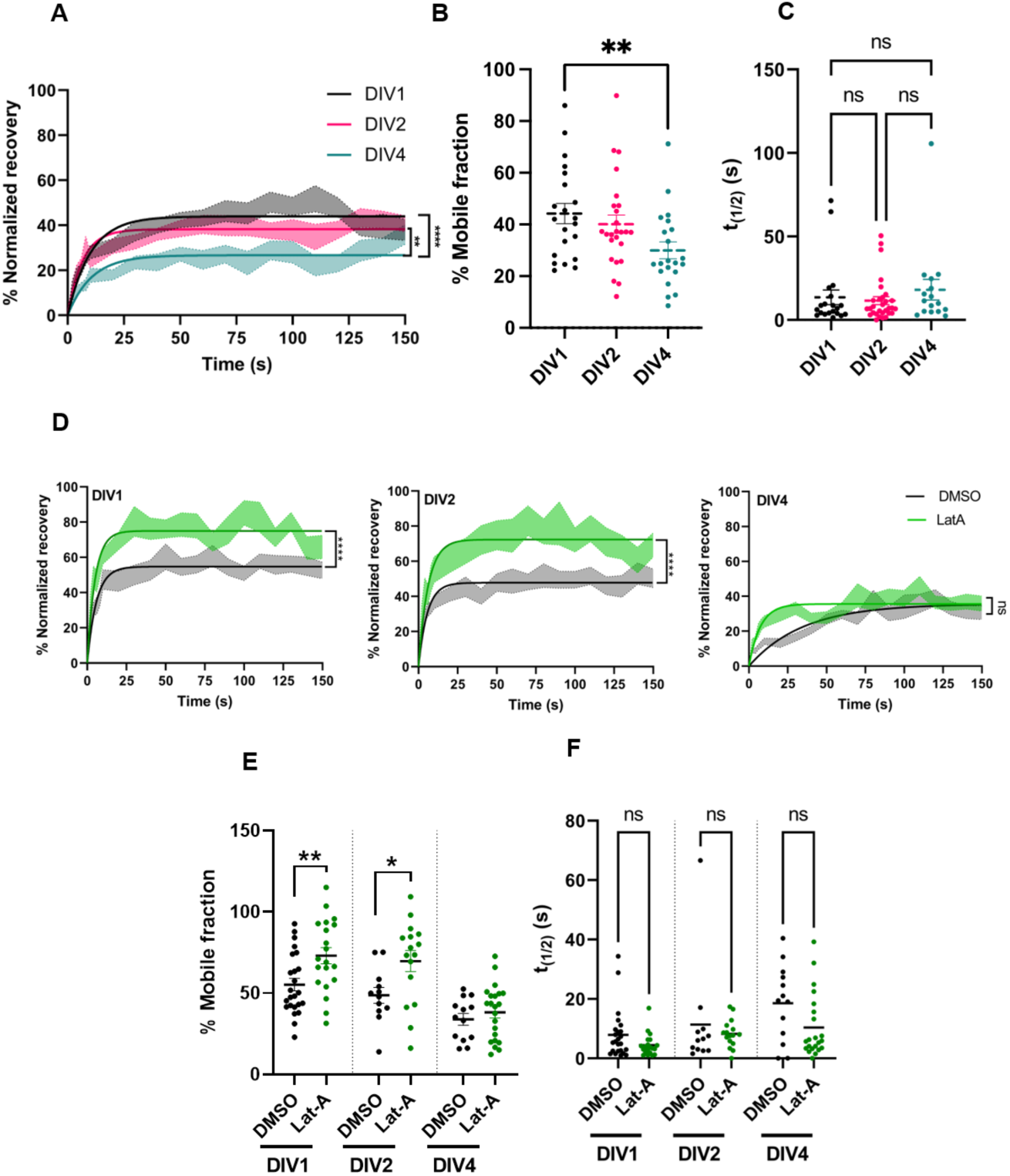
Spectrin mobility is correlated with MPS maturation and dependent on F-actin-mediated cortical recruitment. (A) βII-spectrin-GFP FRAP recovery curves for the DIV1, DIV2, and DIV4 axons. The recovery curves are represented as a one-phase fit of the means (± SEM). Plateau of curves were compared using the extra sum-of-squares F-test. (B) Quantification of the % mobile fraction. The data were analysed using the Mann-Whitney test. (C) *t1/2* values of recovery curves obtained from one-phase fit of Latrunculin A and DMSO-treated DRG neurons. Data were analysed using the Kruskal-Wallis test. For A-C, 16-30 neurons for all DIVs from 3 biological replicates were analysed (D) βII-spectrin-GFP FRAP recovery curves for DRG axons treated with Lat-A across DIV1, DIV2 and DIV4. The green and black traces represent the average one-phase fit for Lat-A- and DMSO-treated neurons, respectively. The recovery curves are represented as a one-phase fit of the means (± SEM). Plateau of curves were compared using the extra sum-of-squares F-test. (E) Quantification of the % mobile fraction. The data were analysed using the Mann-Whitney test. In this experiment, 12-24 neurons for all DIVs from 3 biological replicates were analysed. (F) *t1/2* values of recovery curves obtained from one-phase fit of Latrunculin A and DMSO-treated DRG neurons. Data were analysed using the Kruskal-Wallis test. 12-24 neurons for all DIVs from 3 biological replicates were analysed. (ns, p > 0.05; *, p ≤ 0.05; **, p ≤ 0.01).

Analysis of the normalised recovery curves and the mobile fraction, demonstrated a gradual decrease in the overall mobility of βII-spectrin, with a marked reduction as development progresses from DIV1 to DIV4 (Fig. 6A,B). The half-life (*t_1/2_*) of the recovery was similar across DIVs, indicating no change in the rate of diffusion of βII-spectrin (Fig. 6C). These data are consistent with the increase in the autocorrelation amplitude from DIV1 to DIV4 (Fig. 1), indicating that the βII-spectrin scaffold becomes increasingly stable with development. However, even at the early stages (DIV1), the mobile fraction is under 50%. This suggests that βII-spectrin is already recruited to the axonal cortex by DIV1 with restricted mobility.

Recruitment of spectrin heterotetramers to the cortex may involve interactions with the cortical F-actin. The sensitivity to F-actin deregulation observed earlier (Figs. 2 and 5) likely reflects this association. To explore the role of F-actin in βII-spectrin mobility, we treated βII-spectrin-GFP transfected DIV1, 2, and 4 DRG neurons with 5 μM Lat-A for 15 min (same dose as that used in the data represented in Fig. 2) before FRAP analysis. In contrast to the DMSO-treated control recovery curves for DIV1 and DIV2 neurons, the Lat-A-treated axons exhibited increased βII-spectrin mobility with an elevated mobile fraction (Fig. 6D-E). However, the recovery kinetics of βII-spectrin-GFP in DIV4 axons treated with Lat-A were comparable to DMSO-treated controls. In all cases, there was no change in the *t_1/2_* of the recovery (Fig. 6F).

These data suggest that a significant proportion of βII-spectrin is associated with the axonal cortex as early as DIV1 and constitutes a stable spectrin pool. A major component of this early association is via interactions with subplasmalemmal F-actin, though this dependence is reduced by DIV 4, perhaps due to the recruitment of other interactions that stabilise the association or increased resistance of mature cortical F-actin to depolymerisation. The increased stability of βII-spectrin correlates with the progressive establishment of long-range, periodic order of the MPS with developmental time (Fig. 1).

### F-actin and membrane interactions recruit βII spectrin to the axonal cortex for MPS development

βII-spectrin is recruited to and stably associates with the axonal cortex early in development (Fig. 6). To examine how early recruitment influences MPS development and dissect the underlying mechanisms, we developed differentiated SH-SY5Y human neuroblastoma cells as a model. Using a differentiation protocol (see Methods) involving sequential treatment with retinoic acid (RA) and brain-derived neurotrophic factor (BDNF), we characterised the development of the βII-spectrin MPS in the axon-like SH-SY5Y neurites (Fig. 7A). βII-spectrin autocorrelation and the autocorrelation amplitude both increased following BDNF treatment for 4, 6 and 8 days (Fig. 7B-D). The periodicity measurements demonstrated a progressive decrease in the MPS spacing variability, reaching 191 ± 3.17 nm after 8 days (Fig. 7E), indicating a developmental pattern comparable to DRG neurons (Fig. 1). Next, we generated βII-Spectrin knockout SH-SY5Y cells (βII-Spectrin KO) using CRISPR-Cas9 technology and verified the knockout through genomic sequencing (Fig. S7 A,B) and immunoblotting (Fig. 7 F). As expected, the βII-Spectrin-KO SH-SY5Y neurons displayed reduced axonal lengths (Fig. S7C,D), consistent with the phenotype reported for neurons derived from βII-Spectrin knockout mice (Lorenzo et al., 2019).

**Figure 7:**
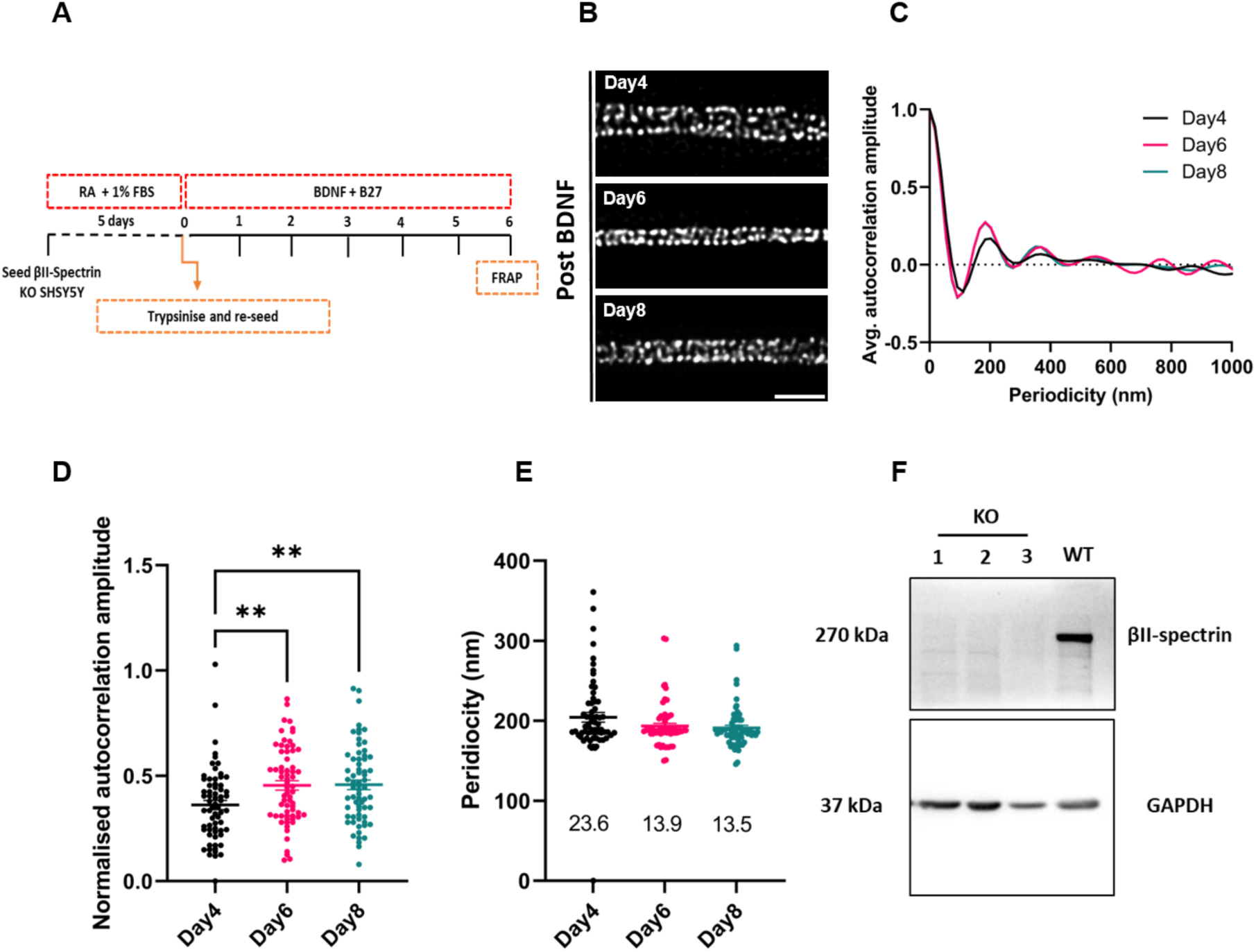
MPS in differentiated SH-SY5Y neurons. (A) Timeline indicating the differentiation protocol of SH-SY5Y cells. (B) Representative micrographs of single optical sections from STED nanoscopy of βII-spectrin stained differentiating neurons (4, 6, and 8 days post-BDNF). Scale bar: 1 µm. (C, D) Average autocorrelation curves and quantification of normalised autocorrelation amplitudes across time points. The data were analysed using the Brown-Forsythe and Welch ANOVA test with multiple comparisons corrected by Games-Howell’s test. (**, p ≤ 0.01). (E) Periodicity across days demonstrates a reduction in variability. The numbers below each data set indicate the coefficient of variation of the distribution. For B-E, 65 axonal segments from 3 biological replicates for each time-point were analysed. (F) Immunoblot validating the CRISPR-Cas9-based generation of βII-spectrin knockout line in SH-SY5Y cells (βII-Spectrin KO). KO-1 was used for all subsequent analyses.

To parse the contributions from various βII-spectrin interactions, we generated variants of βII-spectrin fused with green fluorescent protein (GFP; Fig. 8A). These included the full-length human βII-Spectrin protein, as well as mutants lacking ankyrin B binding (βII-Spectrin-Y1874A), plasma membrane recruitment via the PH domain (βII-Spectrin-K2207Q), and actin-binding (βII-Spectrin-ΔABD). We imaged GFP-labelled axons using STED nanoscopy 6 days after the transfection of these constructs in the βII-Spectrin knockout SH-SY5Y cells (Fig. 8A). The full-length βII-spectrin and ankyrin B binding deficient protein (βII-Spectrin-Y1874A) were recruited to the axonal cortex and formed a periodic scaffold (Fig. 8B-D). However, both the PH domain (βII-Spectrin-K2207Q) and actin-binding domain (βII-Spectrin-ΔABD) mutants were unable to form a strongly periodic MPS, as quantified by the average autocorrelation curves and autocorrelation amplitudes (Fig. 8B-D). Additionally, periodicity analysis showed a higher coefficient of variation for both these mutants compared to the full-length control (Fig. 8E). These findings demonstrate βII-spectrin recruitment to the subplasmalemmal cortex via actin-binding and membrane interactions of the PH-domain to form the MPS.

**Figure 8:**
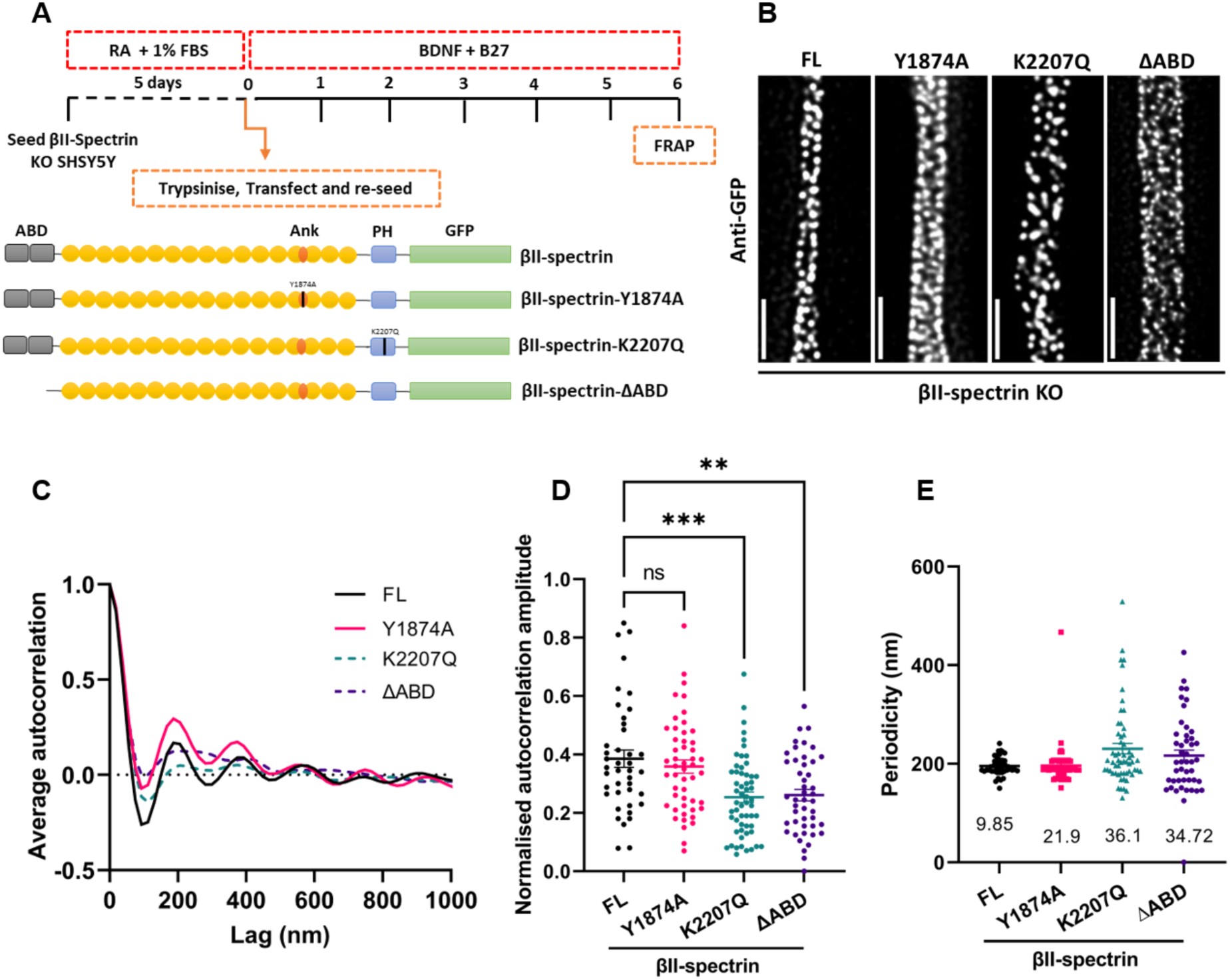
Actin-binding and plasma membrane interactions of βII-spectrin are necessary for MPS development. (A) Timeline indicating the differentiation protocol of βII-Spectrin KO cells and experimental procedures. βII-spectrin KO were transfected mid-differentiation with βII-spectrin variants indicated. The domains are denoted as follows: GFP-tag in green, actin-binding domain (ABD) in grey, ankyrin domain-binding domain in orange, and plasma membrane-binding domain in blue. (B) Representative micrographs of single optical sections from STED nanoscopy of anti-GFP stained differentiated neurites (day 6 post-BDNF) transfected with βII-spectrin variants. (C) Average autocorrelation curves and quantification of normalised autocorrelation amplitudes for all βII-spectrin variants. The data were analysed using the Brown-Forsythe and Welch ANOVA test with multiple comparisons corrected by Games-Howell’s test. (E) Periodicity analysis demonstrates high variation in the spacing between the βII-spectrin rings in K227Q and ΔABD variants. The numbers below each data set indicate the coefficient of variation of the distribution. In this experiment, 40–60 axonal regions were examined for each condition across 3 biological replicates. (ns, p > 0.05; **, p ≤ 0.01; ***, p ≤ 0.001).

To investigate the subplasmalemmal recruitment of βII-spectrin further, we performed FRAP analysis of βII-Spectrin KO cells following transfection with different βII-Spectrin mutants. The recovery kinetics of ankyrin B -binding deficient protein (βII-Spectrin-Y1874A) were comparable to the full-length protein (Fig. 9A,B), suggesting that ankyrin B binding is dispensable at these early stages. However, βII-Spectrin deficient in F-actin binding (βII-Spectrin-ΔABD) had a significantly higher mobile fraction and reduced *t_1/2_* indicating compromised recruitment to the axonal cortex (Fig. 9A,B). This observation is consistent with the elevated mobile fraction of βII-Spectrin observed upon F-actin disruption by Lat-A treatment in primary DRG neurons (Fig. 6D,E).

**Figure 9:**
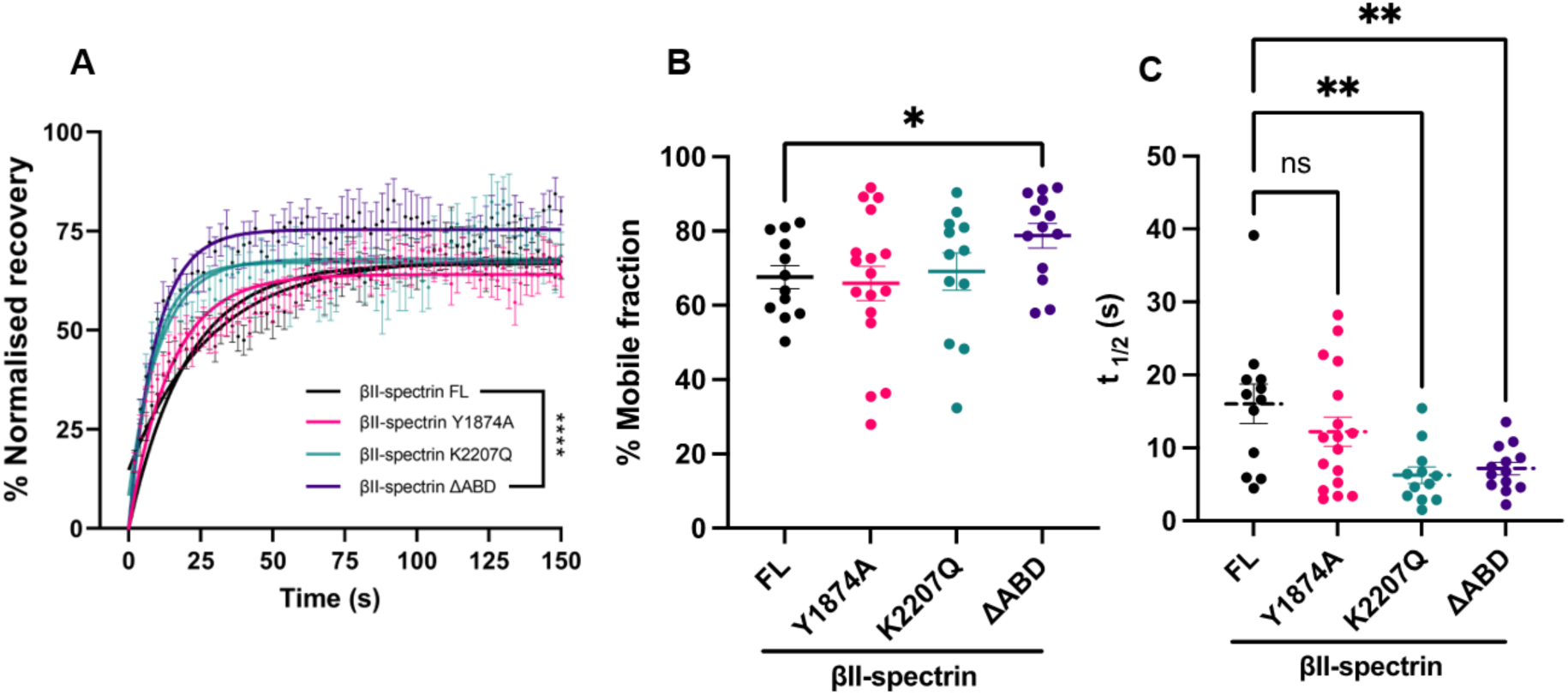
Actin-binding and membrane interactions drive βII-spectrin recruitment and mobility. (A) FRAP recovery curves of mutant and βII-FL-spectrin expressing βII-Spectrin KO cells 6 days after transfection and BDNF treatment. The recovery curves are represented as a one-phase fit of the means (± SEM). Plateau of curves were compared using the extra sum-of-squares F-test. (B) Quantification of the % mobile fraction. The data were analysed using the Mann-Whitney test. (C) t1/2 values extracted from the fit were analysed using the one-way ANOVA test. In this experiment, 12-17 neurons for all DIVs from 3 biological replicates were analysed (ns, p > 0.05; *, p ≤ 0.05; **, p ≤ 0.01; ****, p ≤ 0.0001).

The mobile fraction estimated from the recovery curves of the PH domain mutant(βII-Spectrin-K2207Q) was comparable to the wild-type protein (Fig. 9A,B). However, the *t_1/2_* of the recovery was significantly reduced (Fig. 9C). This suggests no change in the proportion of bound βII-Spectrin-K2207Q molecules but a higher diffusion rate. Given that transfection of βII-Spectrin-K2207Q results in compromised recovery of the MPS (Fig. 8 B-E), this implicates βII-Spectrin PH domain interactions in constraining the kinetics of βII-Spectrin and, in turn, its recruitment to the cortical scaffold.

Collectively, these data identify βII-Spectrin interactions with F-actin and the plasma membrane for early recruitment and stabilisation of βII-Spectrin at the axonal cortex as a necessary prerequisite for MPS development.

## DISCUSSION

Spectrin heterotetramer-based subplasmalemmal scaffolds are common across cell types (Ghisleni et al., 2020; Leterrier and Pullarkat, 2022) but form a unique periodic structure with long-range order in neuronal axons. Given the myriad functions ascribed to the axonal MPS (Albrecht et al., 2016; Lorenzo et al., 2019; Zhou et al., 2019, 2022; Costa et al., 2020), there is considerable interest in understanding the early development of the axonal MPS. Here, we focus on embryonic DRG neurons to uncover the cytoskeleton-mediated mechanisms underlying the recruitment of spectrin to the axonal cortex, the maturation of the βII-Spectrin scaffold to a lattice-like periodic organisation and the maintenance of the MPS.

In this study, we evaluate the medial axonal segments of DRG axons to investigate the establishment of long-range periodic order and the typical spacing of the βII-Spectrin MPS. We find that βII-Spectrin is already present at the axonal cortex by DIV1, though its organisation is weakly periodic. The scaffold matures progressively with development, with the degree of periodicity increasing and the variability of the spectrin spacing concomitantly decreasing and converging to ∼185 nm by DIV 6 (Fig 1A-E). However, there is no obvious change in the amount of βII-Spectrin (Fig S1A) between DIVs indicating re-modelling of early recruited spectrin to a periodic organisation with long-range order. These data also indicate a more rapid development of the MPS in embryonic DRG neurons compared to hippocampal (Zhong et al., 2014) and cortical neurons (Hofmann et al., 2022).

The progressive maturation of the MPS was coincident with a reduction in the axonal diameter. The role of the MPS in regulating axonal diameter has been demonstrated previously (Costa et al., 2020; Wang et al., 2020), and our data suggests maturation of the MPS results in increased radial constriction, possibly due to the increased recruitment of myosin and stabilisation of the F-actin rings. Our data suggest that the corseting of the axon shaft by the MPS regulates axon diameter, and all manipulations that disrupt the MPS result in increased βII-Spectrin ring diameter.These data not only support the correspondence between long-range, periodic MPS and axonal morphology but also indicate that the MPS provides both passive and active (via actomyosin contractility) regulation of the axonal diameter.

Axonal pre-tension also increases modestly with the increase in the periodic organisation, though it is interesting to note that even at DIV1 there is substantial pre-tension. Axonal pre-tension has been previously associated with spectrin (Krieg et al., 2014), and the non-periodic organisation of spectrin in non-neuronal cells are also load-bearing (Lee and Discher, 2001; Smith et al., 2018; Ghisleni et al., 2020). The weakly periodic spectrin organisation observed at DIV1 may form a networked scaffold that supports the axonal pre-stress. However, cytoskeletal components are also likely to contribute significantly as load-bearing structures in the axon (Mutalik et al., 2018; Leterrier and Pullarkat, 2022; Ghose and Pullarkat, 2023).

Acute depolymerisation of microtubules led to the destabilisation of the MPS at all developmental stages (Fig. 2). This suggests a microtubule-dependent MPS maintenance function. Similar observations have been made in hippocampal neurons (Zhong et al., 2014). Conversely, in *Drosophila* neurons, periodic actin is proposed to maintain microtubule alignment and stability (Qu et al., 2017). In *C. elegans*, loss of spectrin reduces the length of axonal microtubules in some neurons (Krieg et al., 2017). To probe the contribution of microtubule to MPS formation further, we subjected early axons to a low chronic dose of microtubule modifying drugs (Fig. 3). As with the acute treatment, continuous exposure to a low dose of Noco from 16h after seeding, resulted in compromised MPS in both DIV 2 and DIV4. Surprisingly, and in contrast to acute treatment with Taxol, chronic stabilisation of microtubules in early axons also destabilised the MPS at both DIV 2 and 4. These observations suggest the requirement of microtubule dynamics in the establishment of the βII-Spectrin MPS. Apart from stabilising microtubules, Taxol can also promote microtubule growth and nucleation, these changes may influence MPS development (Witte et al., 2008; Verma et al., 2016). In epithelial lateral membranes, the mobility of spectrin domains is sensitive to microtubule depolymerisation (Jenkins et al., 2015), however, it is unclear if an analogous or distinct mechanism is involved in axonal MPS development. Recent *in vivo* studies in *C. elegans* suggest a balance between spectrin transport and assembly (Glomb et al., 2023). Though chronic treatment with Noco or Taxol resulted in compromised MPS without any changes in axonal βII-Spectrin intensity, subtle deficits in transport are likely to be unresolved due to the high abundance of relatively stationary spectrin.

Acute depolymerisation but not stabilisation of F-actin resulted in the loss of βII-spectrin MPS across all stages of early development as has also been demonstrated in both young and older axons (Zhong et al., 2014; Han et al., 2017; Qu et al., 2017). A key role of F-actin, in the form of the circumferential rings, involves the crosslinking of spectrin heterotetramers into the longitudinal periodic arrangement seen in axons (Xu et al., 2013; Vassilopoulos et al., 2019). Further, subplasmalemmal F-actin is likely to facilitate the cortical recruitment of spectrin heterotetramers early in development and initiate the organisation of the crosslinked spectrin scaffold.

Inhibition of actin nucleation early in development (DIV2, 4) resulted in disruption of the MPS. However, acute inhibition of actin nucleation at later stages (DIV 6) did not affect the established MPS. This suggests that dynamically nucleating actin networks in early development are required for MPS development, but the contribution of fresh actin nucleation-dependent turnover, over the timescales evaluated, is not necessary for MPS maintenance.

FRAP experiments revealed that βII-spectrin gets increasingly stabilised at the axonal cortex with development, consistent with previous observations in hippocampal neurons (Xu et al., 2013). We show that a significant component of the cortical recruitment of βII-spectrin is via interactions with F-actin, though this dependency reduces as the MPS matures. As βII-spectrin but not αII-spectrin can directly bind F-actin (Li and Bennett, 1996), the latter is expected to drive the interactions of the spectrin heterotetramer with cortical actin.

To further probe subplasmalemmal recruitment of βII-spectrin, we developed a model of differentiated human SH-SY5Y cells engineered to lack βII-spectrin. By expressing βII-spectrin spectrin variants in this system, we established the essential roles of the membrane-interacting PH domain and the actin-binding domain in the development of the MPS. These data are consistent with observations in mouse neurons (Zhou et al., 2022). FRAP analysis of mutant spectrin revealed that the actin-binding domain of βII-spectrin is required for its recruitment and stabilisation at the axonal cortex. Additionally, interactions with the plasma membrane lipids via the PH domain also contribute to subplasmalemmal confinement.

In fibroblasts, actomyosin activity can drive the transition from a diffuse orientation of βII-spectrin to a periodic, clustered configuration (Ghisleni et al., 2024). In contrast, blebbistatin treatment does not affect the MPS organisation in mature neurons, though it increases the axonal diameter by reducing radial contractility (Costa et al., 2020; Wang et al., 2020). However, as early DRG axons are actively contractile (Mutalik et al., 2018), there may be a role for actomyosin contractility in driving the reorganisation of βII-spectrin to a periodic scaffold. Early and chronic inhibition of myosin did not affect axonal MPS formation while increasing the axonal diameter. These data suggest that, unlike in fibroblasts, actomyosin contractility is dispensable for axonal βII-Spectrin MPS development, perhaps due to differences in topological and mechanical constraints.

Recent work has suggested a regulatory coupling between spectrin transport and MPS assembly (Glomb et al., 2023), and there are observations indicative of intercalated MPS assembly in the distal axon (Hofmann et al., 2022; Boyer et al., 2024). Our study focusses on the medial axonal segments and evaluates the developmental remodelling of βII-Spectrin after subplasmalemmal recruitment. Mechanisms identified here are likely to be equally relevant for the distal ends of long axons following cortical recruitment of βII-Spectrin.

This study highlights the dynamic remodelling and maturation of the βII-spectrin MPS in early development and identifies stage-specific cytoskeletal contributions. Dynamics of the microtubule and the actin cytoskeletons contribute to the development of the MPS. Interactions with F-actin and the membrane drive the early subplasmalemmal confinement of βII-spectrin, which subsequently matures to an extended scaffold with long-range periodic order and whose maintenance is dependent on both stable F-actin and microtubules.

## AUTHOR CONTRIBUTIONS

Conceptualisation: S.B. and A.G.; Investigation and formal analysis (all experiments other than laser ablation experiments): S.B.; Investigation and formal analysis (laser ablation experiments): S.B. and A.M.; Methodology and resources: S.B., A.M., P.P. and A.G.; Writing – original draft: S.B. and A.G.; Writing – review and editing: S.B., A.M., P.P. and A.G.; Funding Acquisition: A.G., P.P.

All authors gave final approval for publication and agreed to be held accountable for the work performed therein.

## FUNDING

This work was supported by grant no.: IA/TSG/20/1/600137 from DBT-Wellcome Trust India Alliance, grant no.: DST/INT/Portugal/P-04/2021(G) from the Dept. of Science & Technology, Govt. on India and intramural funds from IISER Pune.

## ACKNOWLEDGEMENTS

The IISER Pune Microscopy Facility is acknowledged for access to imaging systems and software. We thank Dr. Sayantan Majumdar, RRI Bengaluru, for providing the high-speed camera used in the laser ablation experiments.

## Notes

### Competing Interest Statement

The authors have declared no competing interest.

### Summary of Updates

New experiments that augment the conclusion have been included, and the title has been shortened.

## REFERENCES

Albrecht, D., C.M. Winterflood, M. Sadeghi, T. Tschager, F. Noé, and H. Ewers. 2016. Nanoscopic compartmentalization of membrane protein motion at the axon initial segment. J. Cell Biol. 215:37–46. doi:10.1083/jcb.201603108.

Boyer, N.P., R. Sharma, T. Wiesner, A. Delamare, F. Pelletier, C. Leterrier, and S. Roy. 2024. Spectrin condensates provide a nidus for assembling the periodic axonal structure. 2024.06.05.597638. doi:10.1101/2024.06.05.597638.

Bubb, M.R., I. Spector, B.B. Beyer, and K.M. Fosen. 2000. Effects of Jasplakinolide on the Kinetics of Actin Polymerization: AN EXPLANATION FOR CERTAIN IN VIVO OBSERVATIONS*. J. Biol. Chem. 275:5163–5170. doi:10.1074/jbc.275.7.5163.

Costa, A.R., S.C. Sousa, R. Pinto-Costa, J.C. Mateus, C.D. Lopes, A.C. Costa, D. Rosa, D. Machado, L. Pajuelo, X. Wang, F. Zhou, A.J. Pereira, P. Sampaio, B.Y. Rubinstein, I.M. Pinto, M. Lampe, P. Aguiar, and M.M. Sousa. 2020. The membrane periodic skeleton is an actomyosin network that regulates axonal diameter and conduction. eLife. doi:10.7554/eLife.55471.

Cousin, M.A., B.A. Creighton, K.A. Breau, R.C. Spillmann, E. Torti, S. Dontu, S. Tripathi, D. Ajit, R.J. Edwards, S. Afriyie, J.C. Bay, K.M. Harper, A.A. Beltran, L.J. Munoz, L. Falcon Rodriguez, M.C. Stankewich, R.E. Person, Y. Si, E.A. Normand, A. Blevins, A.S. May, L. Bier, V. Aggarwal, G.M.S. Mancini, M.A. van Slegtenhorst, K. Cremer, J. Becker, H. Engels, S. Aretz, J.J. MacKenzie, E. Brilstra, K.L.I. van Gassen, R.H. van Jaarsveld, R. Oegema, G.M. Parsons, P. Mark, I. Helbig, S.E. McKeown, R. Stratton, B. Cogne, B. Isidor, P. Cacheiro, D. Smedley, H.V. Firth, T. Bierhals, K. Kloth, D. Weiss, C. Fairley, J.T. Shieh, A. Kritzer, P. Jayakar, E. Kurtz-Nelson, R.A. Bernier, T. Wang, E.E. Eichler, I.M.B.H. van de Laar, A. McConkie-Rosell, M.T. McDonald, J. Kemppainen, B.C. Lanpher, L.E. Schultz-Rogers, L.B. Gunderson, P.N. Pichurin, G. Yoon, M. Zech, R. Jech, J. Winkelmann, A.S. Beltran, M.T. Zimmermann, B. Temple, S.S. Moy, E.W. Klee, Q.K.-G. Tan, and D.N. Lorenzo. 2021. Pathogenic SPTBN1 variants cause an autosomal dominant neurodevelopmental syndrome. Nat. Genet. 53:1006–1021. doi:10.1038/s41588-021-00886-z.

D’Este, E., D. Kamin, C. Velte, F. Göttfert, M. Simons, and S.W. Hell. 2016. Subcortical cytoskeleton periodicity throughout the nervous system. Sci. Rep. 6:22741. doi:10.1038/srep22741.

Dubey, S., N. Bhembre, S. Bodas, S. Veer, A. Ghose, A. Callan-Jones, and P. Pullarkat. 2020. The axonal actin-spectrin lattice acts as a tension buffering shock absorber. eLife. 9:e51772. doi:10.7554/eLife.51772.

Gallo, G., H.F. Yee Jr., and P.C. Letourneau. 2002. Actin turnover is required to prevent axon retraction driven by endogenous actomyosin contractility. J. Cell Biol. 158:1219–1228. doi:10.1083/jcb.200204140.

Ghisleni, A., M. Bonilla-Quintana, M. Crestani, Z. Lavagnino, C. Galli, P. Rangamani, and N.C. Gauthier. 2024. Mechanically induced topological transition of spectrin regulates its distribution in the mammalian cell cortex. Nat. Commun. 15:5711. doi:10.1038/s41467-024-49906-6.

Ghisleni, A., C. Galli, P. Monzo, F. Ascione, M.-A. Fardin, G. Scita, Q. Li, P. Maiuri, and N.C. Gauthier. 2020. Complementary mesoscale dynamics of spectrin and acto-myosin shape membrane territories during mechanoresponse. Nat. Commun. 11:5108. doi:10.1038/s41467-020-18825-7.

Ghose, A., and P. Pullarkat. 2023. The role of mechanics in axonal stability and development. Semin. Cell Dev. Biol. 140:22–34. doi:10.1016/j.semcdb.2022.06.006.

Glomb, O., G. Swaim, P. Munoz LLancao, C. Lovejoy, S. Sutradhar, J. Park, Y. Wu, S.E. Cason, E.L.F. Holzbaur, M. Hammarlund, J. Howard, S.M. Ferguson, M.W. Gramlich, and S. Yogev. 2023. A kinesin-1 adaptor complex controls bimodal slow axonal transport of spectrin in *Caenorhabditis elegans*. Dev. Cell. 58:1847–1863.e12. doi:10.1016/j.devcel.2023.08.031.

Hammarlund, M., E.M. Jorgensen, and M.J. Bastiani. 2007. Axons break in animals lacking β-spectrin. J. Cell Biol. 176:269–275. doi:10.1083/jcb.200611117.

Han, B., R. Zhou, C. Xia, and X. Zhuang. 2017. Structural organization of the actin-spectrin–based membrane skeleton in dendrites and soma of neurons. Proc. Natl. Acad. Sci. 114:E6678–E6685. doi:10.1073/pnas.1705043114.

He, J., R. Zhou, Z. Wu, M.A. Carrasco, P.T. Kurshan, J.E. Farley, D.J. Simon, G. Wang, B. Han, J. Hao, E. Heller, M.R. Freeman, K. Shen, T. Maniatis, M. Tessier-Lavigne, and X. Zhuang. 2016. Prevalent presence of periodic actin–spectrin-based membrane skeleton in a broad range of neuronal cell types and animal species. Proc. Natl. Acad. Sci. 113:6029–6034. doi:10.1073/pnas.1605707113.

Hofmann, M., L. Biller, U. Michel, M. Bähr, and J.C. Koch. 2022. Cytoskeletal assembly in axonal outgrowth and regeneration analyzed on the nanoscale. Sci. Rep. 12:14387. doi:10.1038/s41598-022-18562-5.

Huang, C.Y.-M., C. Zhang, T.S.-Y. Ho, J. Oses-Prieto, A.L. Burlingame, J. Lalonde, J.L. Noebels, C. Leterrier, and M.N. Rasband. 2017a. αII Spectrin Forms a Periodic Cytoskeleton at the Axon Initial Segment and Is Required for Nervous System Function. J. Neurosci. 37:11311–11322. doi:10.1523/JNEUROSCI.2112-17.2017.

Huang, C.Y.-M., C. Zhang, D.R. Zollinger, C. Leterrier, and M.N. Rasband. 2017b. An αII Spectrin-Based Cytoskeleton Protects Large-Diameter Myelinated Axons from Degeneration. J. Neurosci. 37:11323–11334. doi:10.1523/JNEUROSCI.2113-17.2017.

Jenkins, P.M., M. He, and V. Bennett. 2015. Dynamic spectrin/ankyrin-G microdomains promote lateral membrane assembly by opposing endocytosis. Sci. Adv. 1:e1500301. doi:10.1126/sciadv.1500301.

Krieg, M., A.R. Dunn, and M.B. Goodman. 2014. Mechanical control of the sense of touch by β-spectrin. Nat. Cell Biol. 16:224–233. doi:10.1038/ncb2915.

Krieg, M., J. Stühmer, J.G. Cueva, R. Fetter, K. Spilker, D. Cremers, K. Shen, A.R. Dunn, and M.B. Goodman. 2017. Genetic defects in β-spectrin and tau sensitize C. elegans axons to movement-induced damage via torque-tension coupling. eLife. 6:e20172. doi:10.7554/eLife.20172.

Lee, J.C.-M., and D.E. Discher. 2001. Deformation-Enhanced Fluctuations in the Red Cell Skeleton with Theoretical Relations to Elasticity, Connectivity, and Spectrin Unfolding. Biophys. J. 81:3178–3192. doi:10.1016/S0006-3495(01)75954-1.

Leterrier, C., and P.A. Pullarkat. 2022. Mechanical role of the submembrane spectrin scaffold in red blood cells and neurons. J. Cell Sci. 135:jcs259356. doi:10.1242/jcs.259356.

Li, X., and V. Bennett. 1996. Identification of the Spectrin Subunit and Domains Required for Formation of Spectrin/Adducin/Actin Complexes *. J. Biol. Chem. 271:15695–15702. doi:10.1074/jbc.271.26.15695.

Lorenzo, D.N., A. Badea, R. Zhou, P.J. Mohler, X. Zhuang, and V. Bennett. 2019. βII-spectrin promotes mouse brain connectivity through stabilizing axonal plasma membranes and enabling axonal organelle transport. doi:10.1073/pnas.1820649116.

Mutalik, S.P., J. Joseph, P.A. Pullarkat, and A. Ghose. 2018. Cytoskeletal Mechanisms of Axonal Contractility. Biophys. J. 115:713–724. doi:10.1016/j.bpj.2018.07.007.

Nishimura, Y., S. Shi, F. Zhang, R. Liu, Y. Takagi, A.D. Bershadsky, V. Viasnoff, and J.R. Sellers. 2021. The formin inhibitor SMIFH2 inhibits members of the myosin superfamily. J. Cell Sci. 134:jcs253708. doi:10.1242/jcs.253708.

Qu, Y., I. Hahn, S.E.D. Webb, S.P. Pearce, and A. Prokop. 2017. Periodic actin structures in neuronal axons are required to maintain microtubules. Mol. Biol. Cell. 28:296–308. doi:10.1091/mbc.e16-10-0727.

Ran, F.A., P.D. Hsu, J. Wright, V. Agarwala, D.A. Scott, and F. Zhang. 2013. Genome engineering using the CRISPR-Cas9 system. Nat. Protoc. 8:2281–2308. doi:10.1038/nprot.2013.143.

Smith, A.S., R.B. Nowak, S. Zhou, M. Giannetto, D.S. Gokhin, J. Papoin, I.C. Ghiran, L. Blanc, J. Wan, and V.M. Fowler. 2018. Myosin IIA interacts with the spectrin-actin membrane skeleton to control red blood cell membrane curvature and deformability. Proc. Natl. Acad. Sci. 115:E4377–E4385. doi:10.1073/pnas.1718285115.

Unsain, N., M.D. Bordenave, G.F. Martinez, S. Jalil, C. von Bilderling, F.M. Barabas, L.A. Masullo, A.D. Johnstone, P.A. Barker, M. Bisbal, F.D. Stefani, and A.O. Cáceres. 2018. Remodeling of the Actin/Spectrin Membrane-associated Periodic Skeleton, Growth Cone Collapse and F-Actin Decrease during Axonal Degeneration. Sci. Rep. 8:3007. doi:10.1038/s41598-018-21232-0.

Vassilopoulos, S., S. Gibaud, A. Jimenez, G. Caillol, and C. Leterrier. 2019. Ultrastructure of the axonal periodic scaffold reveals a braid-like organization of actin rings. Nat. Commun. 10:5803. doi:10.1038/s41467-019-13835-6.

Verma, S., N. Kumar, and V. Verma. 2016. Role of paclitaxel on critical nucleation concentration of tubulin and its effects thereof. Biochem. Biophys. Res. Commun. 478:1350–1354. doi:10.1016/j.bbrc.2016.08.127.

Wang, G., D.J. Simon, Z. Wu, D.M. Belsky, E. Heller, M.K. O’Rourke, N.T. Hertz, H. Molina, G. Zhong, M. Tessier-Lavigne, and X. Zhuang. 2019. Structural plasticity of actin-spectrin membrane skeleton and functional role of actin and spectrin in axon degeneration. eLife. 8:e38730. doi:10.7554/eLife.38730.

Wang, T., W. Li, S. Martin, A. Papadopulos, M. Joensuu, C. Liu, A. Jiang, G. Shamsollahi, R. Amor, V. Lanoue, P. Padmanabhan, and F.A. Meunier. 2020. Radial contractility of actomyosin rings facilitates axonal trafficking and structural stability. J. Cell Biol. 219:e201902001. doi:10.1083/jcb.201902001.

Wernert, F., S.B. Moparthi, F. Pelletier, J. Lainé, E. Simons, G. Moulay, F. Rueda, N. Jullien, S. Benkhelifa-Ziyyat, M.-J. Papandréou, C. Leterrier, and S. Vassilopoulos. 2024. The actin-spectrin submembrane scaffold restricts endocytosis along proximal axons. Science. 385:eado2032. doi:10.1126/science.ado2032.

Witte, H., D. Neukirchen, and F. Bradke. 2008. Microtubule stabilization specifies initial neuronal polarization. J. Cell Biol. 180:619–632. doi:10.1083/jcb.200707042.

Xu, K., G. Zhong, and X. Zhuang. 2013. Actin, Spectrin, and Associated Proteins Form a Periodic Cytoskeletal Structure in Axons. Science. 339:452–456. doi:10.1126/science.1232251.

Zhong, G., J. He, R. Zhou, D. Lorenzo, H.P. Babcock, V. Bennett, and X. Zhuang. 2014. Developmental mechanism of the periodic membrane skeleton in axons. eLife. 3:e04581. doi:10.7554/eLife.04581.

Zhou, R., B. Han, R. Nowak, Y. Lu, E. Heller, C. Xia, A.H. Chishti, V.M. Fowler, and X. Zhuang. 2022. Proteomic and functional analyses of the periodic membrane skeleton in neurons. Nat. Commun. 13:3196. doi:10.1038/s41467-022-30720-x.

Zhou, R., B. Han, C. Xia, and X. Zhuang. 2019. Membrane-associated periodic skeleton is a signaling platform for RTK transactivation in neurons. Science. 365:929–934. doi:10.1126/science.aaw5937.

