## Supplementary Figures for "Development of the axonal βII-spectrin periodic skeleton requires active cytoskeletal remodelling"

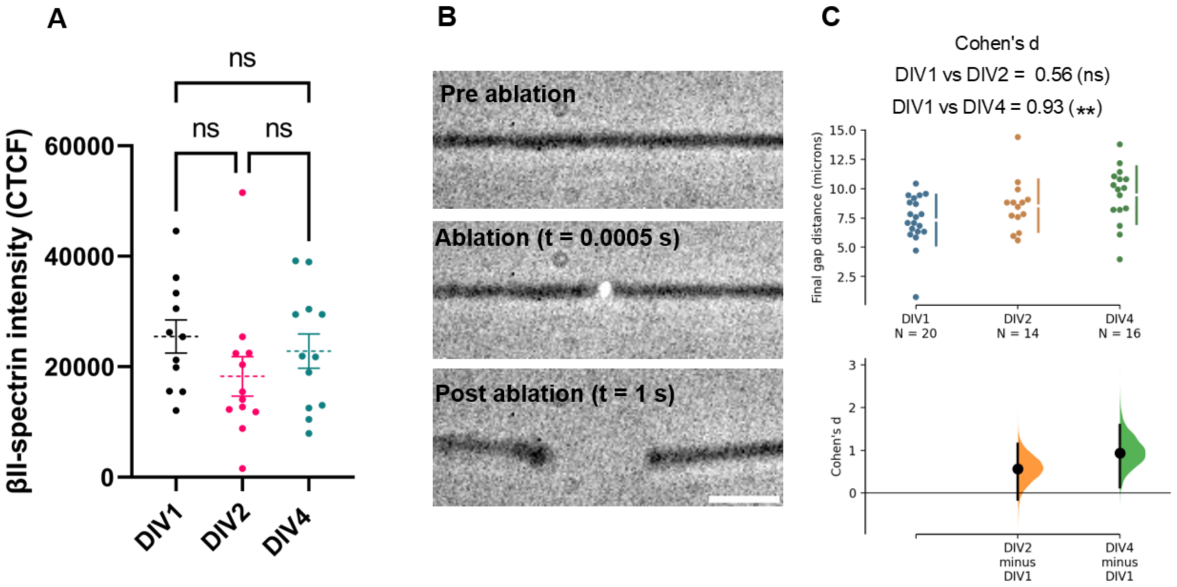


**Figure S1: Developmental time course of βII-spectrin expression and axonal pre-tension.** (A) βII-spectrin intensity in the middle region of the axon is shown as corrected total cell fluorescence (CTCF). The data were analysed using the Kruskal-Wallis test with multiple comparison corrected by Dunn's test (ns, $p > 0.05$). The data are derived from 10-12 axons for all DIVs. (B) Three representative frames from an axonal ablation time series recorded at 2000 fps. Scale bar: 5 μm. (C) Comparison of the final gap distance (at 0.74 sec) post-ablation across DIV1, DIV2 and DIV4 axons was performed using Estimation statistics. The effect size (Cohen's d) and statistical significance (two-sided permutation t-test) are indicated. The data are derived from n=19 (DIV1), n=14 (DIV2) and n=16 (DIV4) axons from three independent biological replicates. The unpaired Cohen's d between DIV1 and DIV2 is 0.564 [95.0%CI -0.151, 1.15] with p = 0.123 (ns) and between DIV1 and DIV4 is 0.934 95.0%CI 0.136, 1.59] with p = 0.0078 (**).

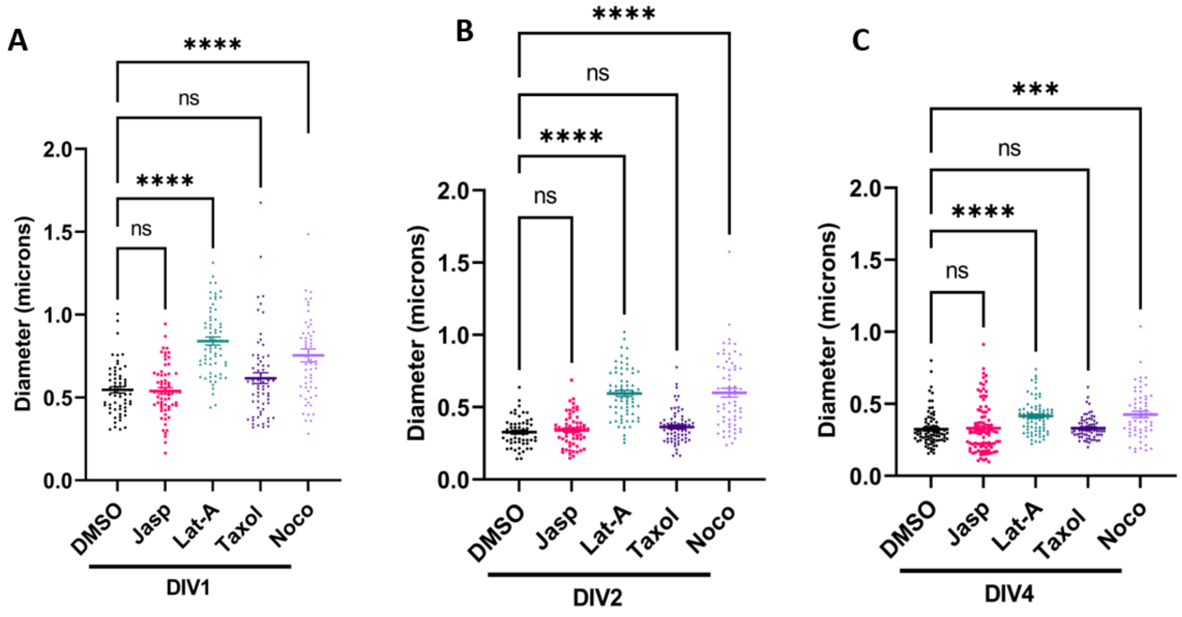


**Figure S2: Microtubule and F-actin destabilisation increases axonal diameter across DIVs.** (A, B, C) Diameter analysis of DMSO, Taxol, Nocodazole and Jasplakinolide-treated axons across DIVs. Data was analysed using the Kruskal-Wallis test with Dunn's correction. 55-95 axonal segments from 3 biological replicates were analysed (ns, $p > 0.05$; ***, $p \le 0.001$ *****, $p \le 0.0001$).

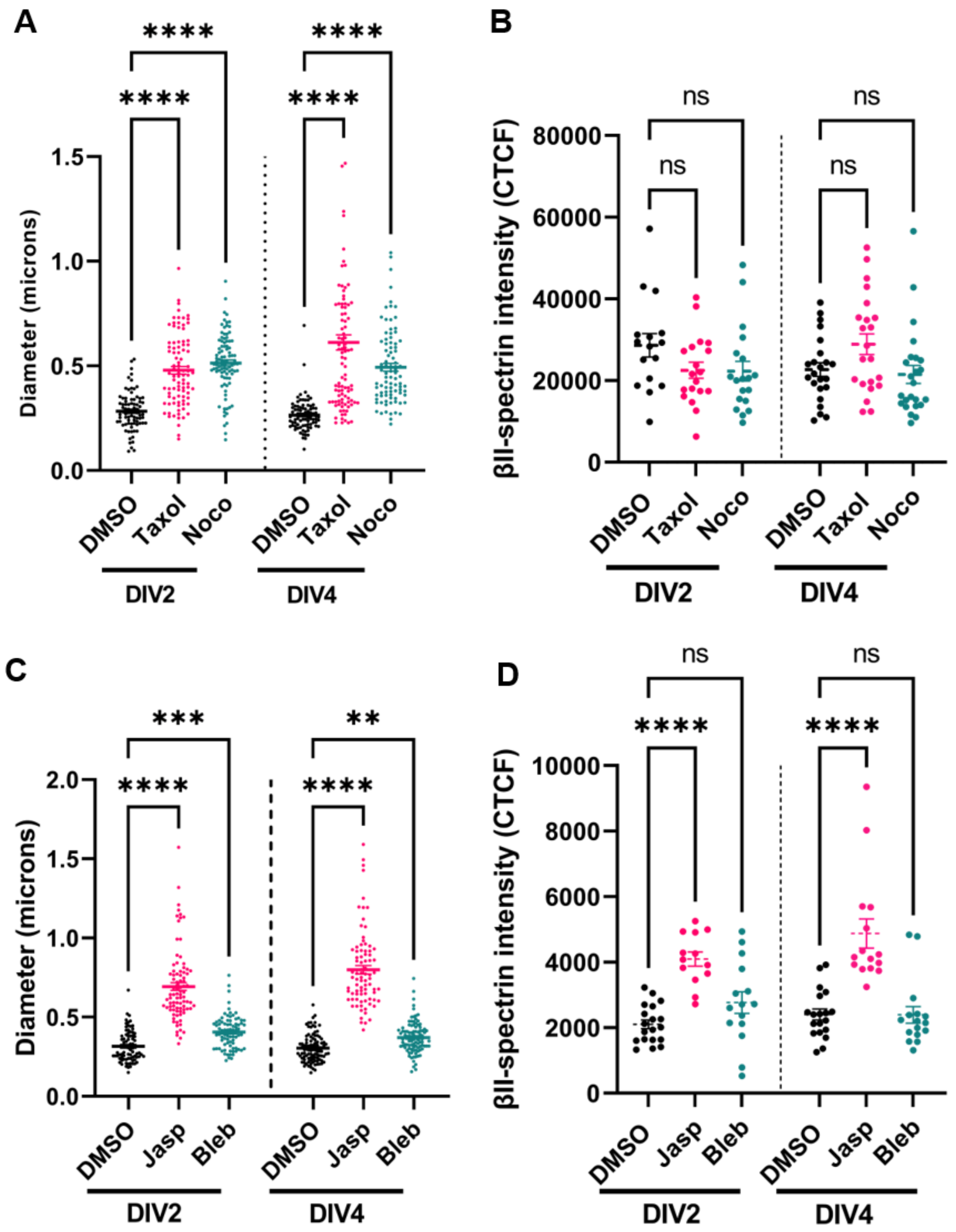


**Figure S3: βII-spectrin ring diameter and βII-spectrin intensity analysis of axons chronically treated with cytoskeleton drugs.** (A) Diameter analysis of DMSO, Taxol and Nocodazole-treated axons across DIVs. Data was analysed using the Kruskal-Wallis test with Dunn's correction. 82-100 axonal segments from 3 biological replicates were analysed **(*******, $p \leq 0.0001$). **(**B) βII-spectrin intensity in the middle region of the axons treated with Taxol and Nocodazole is shown as corrected total cell fluorescence (CTCF). The data were analysed using the Kruskal-Wallis test with multiple comparisons corrected by Dunn's test. The data are derived from 16-24 axons for all DIVs (ns, $p > 0.05$). (C) Diameter analysis of DMSO, Jasplakinolide and Blebbistatin-treated axons across DIVs. Data was analysed using the Kruskal-Wallis test with Dunn's correction. 79-98 axonal segments from 3 biological replicates were analysed. **(**D) βII-spectrin intensity in the middle region of the axons treated with Jasplakinolide and Blebbistatin is shown as corrected total cell fluorescence (CTCF). The data were analysed using the Kruskal-Wallis test with multiple comparisons corrected by Dunn's test. The data are derived from 13-19 axons for all DIVs (ns, $p > 0.05$; **, $p \leq 0.01$; ***, $p \leq 0.001$ *****, $p \leq 0.0001$)**.**

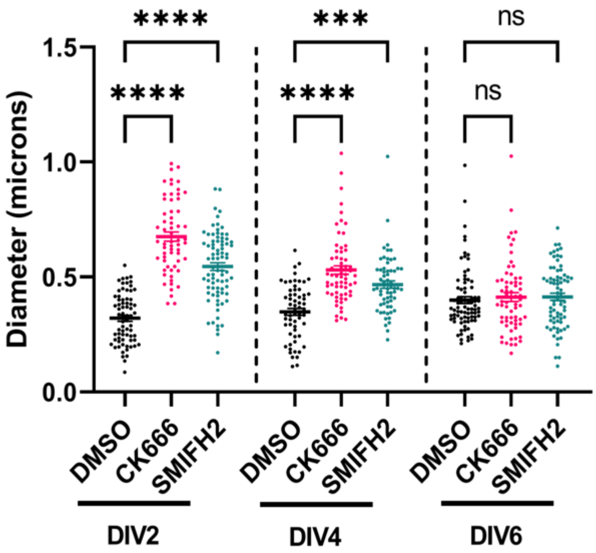


**Figure S4: SMIFH2 or CK666 increases βII-spectrin diameter at DIV2 and DIV4 but not at DIV6.** (A) Diameter analysis of DMSO, SMIFH2- and CK666-treated axons across DIVs. Data was analysed using the Kruskal-Wallis test with Dunn's correction. 60-70 axonal segments from 3 biological replicates were analysed **(**ns, $p > 0.05$; *****, $p \leq 0.0001$).

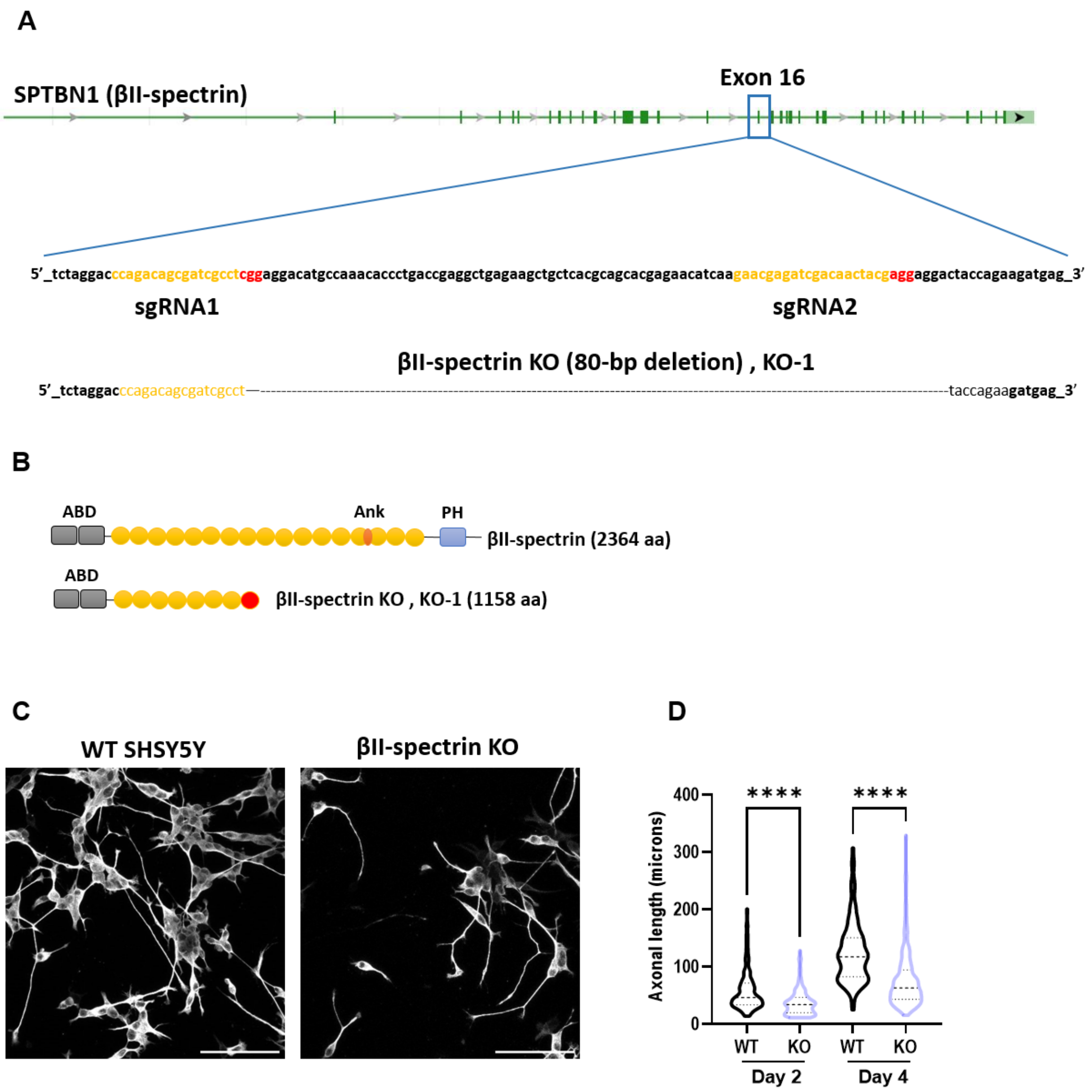


**Figure S5: Generation of βII-spectrin deficient SH-SY5Y cell line.** (A) Sequence analysis of SH-SY5Y βII-spectrin knockout cells (βII-Spectrin KO). Sequences targeted by gRNA 1 and 2 are highlighted in yellow, and the PAM sequences are in red. All subsequent experiments with βII-Spectrin KO use clone KO-1. This clone carries an 80 bp deletion indicated in the sequence. B) Full-length βII-spectrin protein with annotated domains along with the predicted protein in βII-Spectrin KO, clone KO-1. The 80 bp deletion is expected to generate a potential truncated protein (1158 aa) with a premature stop codon highlighted in red. (C) Representative micrographs of differentiated wildtype and βII-Spectrin KO cells stained with βIII tubulin (day 4 post-BDNF). (D) Axonal length quantification by βIII tubulin staining of differentiating wild-type and βII-spectrin knockout SH-SY5Y cells at day 2 and 4 post-BDNF treatment. The data were analysed using the one-way ANOVA; ****, $p \leq 0.0001$. 20-30 neurons from 3 independent biological replicates were analysed for each genotype and days post-BDNF.
